# Proteomic and *in silico* dissection of MetaAggregates in amyotrophic lateral sclerosis brains

**DOI:** 10.1101/2025.07.09.663594

**Authors:** Kazuma Murakami, Thi Hong Van Nguyen, Nobuko Fujita, Chioko Nagao, Kenji Mizuguchi, Takumi Nishiuchi, Yasuhiro Sakashita, Moeko Noguchi-Shinohara, Kenjiro Ono

**Affiliations:** Division of Food Science and Biotechnology, Graduate School of Agriculture, Kyoto University, Kyoto 606-8502, Japan; Institute for Protein Research, The University of Osaka, Osaka 565-0871, Japan; Center for Experimental Modeling of Human Disease, Kanazawa University, 13-1 Takaramachi, Kanazawa, Ishikawa, 920-8640, Japan; Department of Neurology, Kanazawa University Graduate School of Medical Sciences, Kanazawa 920-8640, Japan

**Keywords:** amyotrophic lateral sclerosis, G-quadruplex, RNA-binding protein, single-cell RNA-seq

## Abstract

RNA-binding proteins (RBPs), key translation regulators, are thought to be involved in the pathogenesis of amyotrophic lateral sclerosis (ALS). The pathological entities associated with ALS are known as “MetaAggregates”: heterogeneous coaggregates composed of amyloids, RBPs, and RNA G-quadruplexes (rG4s). In this study, to explore the molecular constituents of ALS-associated MetaAggregates, we developed a proteomic approach using a psoralen-conjugated RBP and crosslinked it with a biotinylated rG4 to enable the isolation of MetaAggregates from ALS brain extracts. Single-cell RNA-seq using *in vitro* ALS models identified ELAVL4 as a cytoplasmic RBP and revealed the enrichment of an IGFBP2-derived rG4 structure in ALS-specific neurons. Mass spectrometry and amyloidogenicity-based principal component analysis revealed 79 candidate proteins with roles in RNA processing, metabolism, trafficking, and stress responses. Docking simulations highlighted a subset of proteins with potential pro-aggregation characteristics, diverse cytosolic associations and functional links to RNA processing relevant to ALS. Through proteomic and *in silico* dissection of ALS-associated MetaAggregates, the findings of this study establish a conceptual framework for the exploration of unrecognized amyloidogenic drivers of neurodegeneration.

## Introduction

Neurodegenerative diseases such as amyotrophic lateral sclerosis (ALS) are characterized by progressive neuronal loss within the central nervous system (CNS), disrupted neural communication, significant morbidity, and often premature mortality. Most neurodegenerative disorders are age-associated, with their central pathological mechanism being the misfolding and aberrant self-assembly of amyloidogenic proteins (Marsh, 2019). To date, more than 50 human amyloidogenic proteins and peptides, collectively referred to as “amyloidoses” or “proteinopathies,” have been identified that either cause or significantly contribute to the onset and progression of these conditions (Bayer, 2015). The CNS is particularly vulnerable to the accumulation of toxic protein aggregates and oligomers due to its high metabolic demands and the inability of neurons to dilute proteinopathies through cell division. Well-known examples of pathogenic proteinaceous species include the tau and amyloid β (Aβ) proteins in Alzheimer’s disease (AD) (Goedert, 2020; Selkoe, 2001) and other tauopathies; α-synuclein (αSyn) in Parkinson’s disease (PD) and other synucleinopathies (Henderson et al, 2019; Spillantini & Goedert, 2000).

These proteins have an inherent tendency to misfold and self-assemble into neurotoxic oligomers and amyloid fibrils (Murakami et al, 2025). *In vitro*, this process often proceeds until most monomers are incorporated into aggregates, followed by a plateau phase with no further growth (Hasegawa et al, 1999; Hellstrand et al, 2010; Ilie & Caflisch, 2019). Importantly, metastable and dynamically fluctuating oligomers have frequently been shown to be more toxic than mature, structurally stable fibrils (Benilova et al, 2012; Bitan et al, 2005; Murakami, 2014; Murakami et al, 2022a; Murakami et al., 2025; Roychaudhuri et al, 2009). These toxicities are mediated through diverse mechanisms, including apoptosis, oxytosis/ferroptosis, chronic inflammation, and disrupted proteostasis. Therefore, oligomeric species have attracted considerable interest as molecular targets for diagnostic and therapeutic strategies.

Meta-aggregates, an emerging concept in proteinopathy, are metastable heterogeneous coaggregates that are formed *in cellulo*. They can consist of combinations of amyloids (e.g., Aβ/αSyn, Aβ/tau, αSyn/tau; (Colom-Cadena et al, 2013; Irwin et al, 2013; Moussaud et al, 2014; Murakami & Ono, 2022; Spires-Jones et al, 2017; Zekry et al, 2002), in which formation they are known as bipartite amyloids, or of amyloids coaggregated with specific biomolecules such as nucleic acids or proteins (**Fig. 1A**). Among the nucleic acid / amyloid complexes, RNA G-quadruplexes (rG4s) are particularly important. These quadruple-stranded structures, stabilized by contiguous guanine tracts, are known to serve as signals for neurite mRNA targeting (Subramanian et al, 2011). Recent studies have shown that rG4s can act as scaffolds to facilitate the aggregation of αSyn (Matsuo et al, 2024) and tau (Yabuki et al, 2024). rG4s undergo Ca²⁺-induced phase separation and assembly, thereby accelerating the sol–gel phase transition of αSyn. In neurons treated with αSyn preformed fibrils, elevated cytoplasmic Ca²⁺ levels have been reported to promote the assembly of rG4s containing synaptic mRNAs, which coaggregate with αSyn and contribute to synaptic dysfunction (Matsuo et al., 2024). These findings highlight the role of rG4s as multifunctional, non-canonical RNA structures that regulate gene expression, RNA stability, and translation, while also playing critical roles in cellular stress responses and disease pathogenesis.

**Figure 1.**
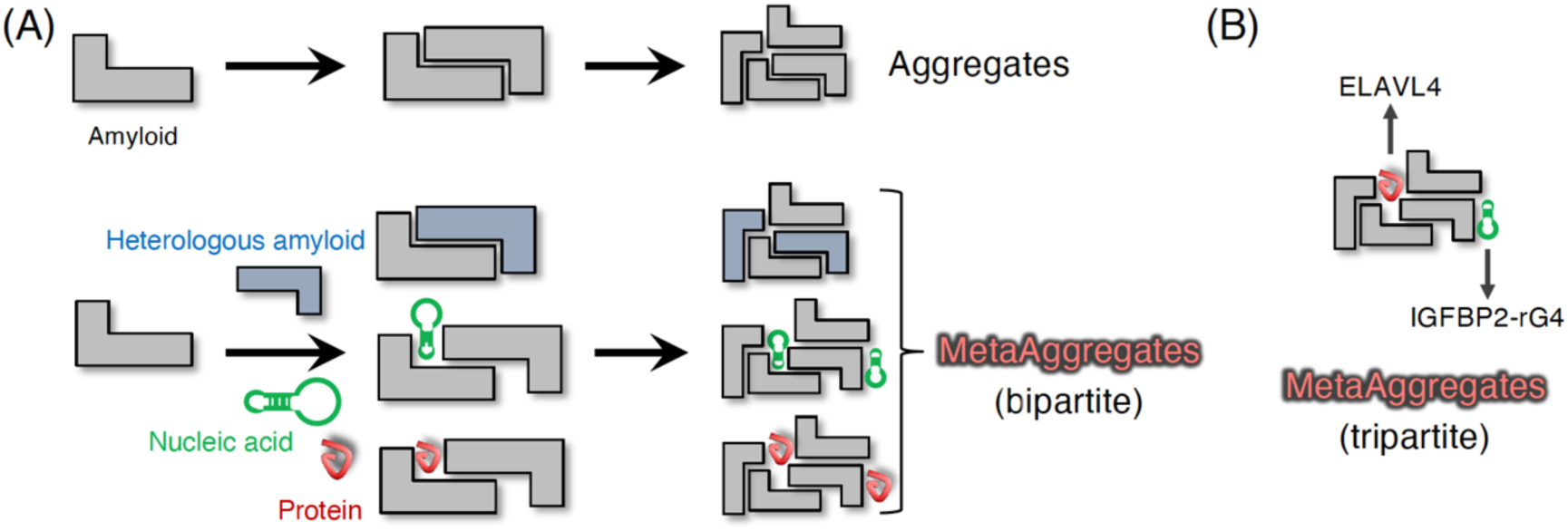
Representative models of metaaggregate formations via coaggregation of amyloids or with non-amyloid biomolecules. **(A)** MetaAggregates are heterogeneous and metastable protein assemblies formed by the coaggregation of distinct amyloid species (e.g., Aβ with αSyn, Aβ with tau, or αSyn with tau), or through interactions with other biomolecules such as nucleic acids and proteins. **(B)** Hypothetical model of tripartite MetaAggregate formation in ALS pathology. In ALS, rG4s, RBP, and amyloids are proposed to coassemble into a tripartite meta-aggregate. For instance, rG4 derived from C9orf72-repeat expansions may recruit ELAVL4, facilitating aberrant aggregation of amyloidogenic proteins, contributing to ALS pathogenesis.

ALS is a fatal neurodegenerative disorder characterized by the progressive loss of motor neurons, leading to muscle weakness, paralysis, and eventual respiratory failure (Brown & Al-Chalabi, 2017; Masrori & Van Damme, 2020). Despite extensive research, the molecular mechanisms underlying the progression of ALS have not been fully elucidated. A key pathological feature of sporadic ALS, which accounts for 90% of cases, is the abnormal aggregation of RNA-binding proteins (RBPs), such as TAR DNA-binding protein 43 (TDP-43) that disrupt RNA metabolism and cellular homeostasis (Mead et al, 2023); this suggests a broader role of RBPs in ALS disease pathology (Arai et al, 2006; Neumann et al, 2006). Similar pathological aggregates are also observed in frontotemporal dementia (FTD), which shares genetic and molecular overlap with ALS. Similarly, TDP-43-negative sporadic ALS brains often exhibit aggregates of SOD1/fused-in-sarcoma (FUS) (Vance et al, 2009) heterogeneous nuclear ribonucleoproteins (hnRNPs) (Gilpin et al, 2015) (Kato et al, 2001), further emphasizing the heterogeneous nature of ALS-associated proteinopathy (Mackenzie et al, 2007). T cell intracellular antigen-1 (TIA-1) plays a central role in stress granule (SG) formation by self-assembly via the prion-like domain, where amino acid mutations associated with ALS and FTD have been identified (Ito et al, 2017; Taylor et al, 2016). These mutations are thought to transform the SG constituents into the aberrant aggregates that accumulate in neurons in these diseases (Wolozin & Ivanov, 2019). Moreover, a hexanucleotide (GGGGCC) repeat expansion in the *C9orf72* gene, the most common genetic cause of ALS and FTD, has been shown to form aberrant RNA foci and dipeptide repeat (DPR) proteins via repeat-associated non-AUG translation (Ramos-Campoy et al, 2018). The DPR proteins, which include poly(GA), poly(GP), and poly(GR), exhibit amyloid-like properties and promote the aggregation of TDP-43, leading to neurotoxicity through mechanisms such as nucleocytoplasmic transport defects, proteasomal inhibition, and phase separation disturbances that exacerbates ALS pathology (Balendra & Isaacs, 2018; Mori et al, 2013). In contrast to AD and PD, which are characterized by specific amyloid pathologies, ALS lacks a single unifying amyloid species across cases; instead, its pathogenesis appears to involve multiple aggregation-prone proteins (Balendra & Isaacs, 2018; Brettschneider et al, 2015; Taylor et al., 2016).

Accumulating evidence has suggested that in ALS pathology RBPs play crucial roles in amyloid aggregation modulation through coaggregation mechanisms. TDP-43, a hallmark RBP in ALS and FTD, has been shown to coaggregate with tau in AD brains (Amador-Ortiz et al, 2007), with tau or αSyn in AD and DLB (dementia with Lewy bodies) brains (Higashi et al, 2007), and with oligomeric Aβ in AD brains (Montalbano et al, 2020). Ash et al. (2021) reported that TIA1, another ALS/FTD-linked RBP, potentiated tau phase separation and induced tau toxic oligomer formation in mouse primary neuronal culture. TIA1 has been also found to coaggregate with tau in transgenic P301S Tau mice (Apicco et al, 2018). These findings underscore the pathological relevance of RBP/amyloid coaggregated bipartite MetaAggregates as a unifying mechanism across diverse neurodegenerative conditions (**Fig. 1A**). In particular, *C9orf72*-originated rG4s interact with RBPs, influencing the formation and aggregation of pathological protein complexes in ALS (Haeusler et al, 2016). This raises the intriguing possibility that RNA, RBPs, and amyloids may coassemble into a tripartite MetaAggregates in ALS pathology (**Fig. 1B**).

In this study, we employed a comprehensive proteomic and bioinformatic approach using single-cell RNA sequencing (scRNA-seq) data from differentiated induced pluripotent stem cells (iPSCs) to search for tripartite MetaAggregates in ALS brains. We characterized ELAV-like protein 4 (ELAVL4) as a key RBP and an RNA oligonucleotide (IGFBP2-rG4) in insulin-like growth factor binding protein 2 (IGFBP2) as an rG4 sequence. To develop a chemical probe to study ALS-associated amyloids, mimetics of tripartite MetaAggregates were synthesized using psoralen and diazirine-mediated photo-crosslinking technology, which enabled the identification of ELAVL4-associated aggregative proteins after incubation with ALS brain lysates (**Fig. 2A**). To determine the impact of structural stability on the roles of MetaAggregates in ALS pathogenesis, we also performed *in silico* docking analyses to explore the structural perspectives.

**Figure 2.**
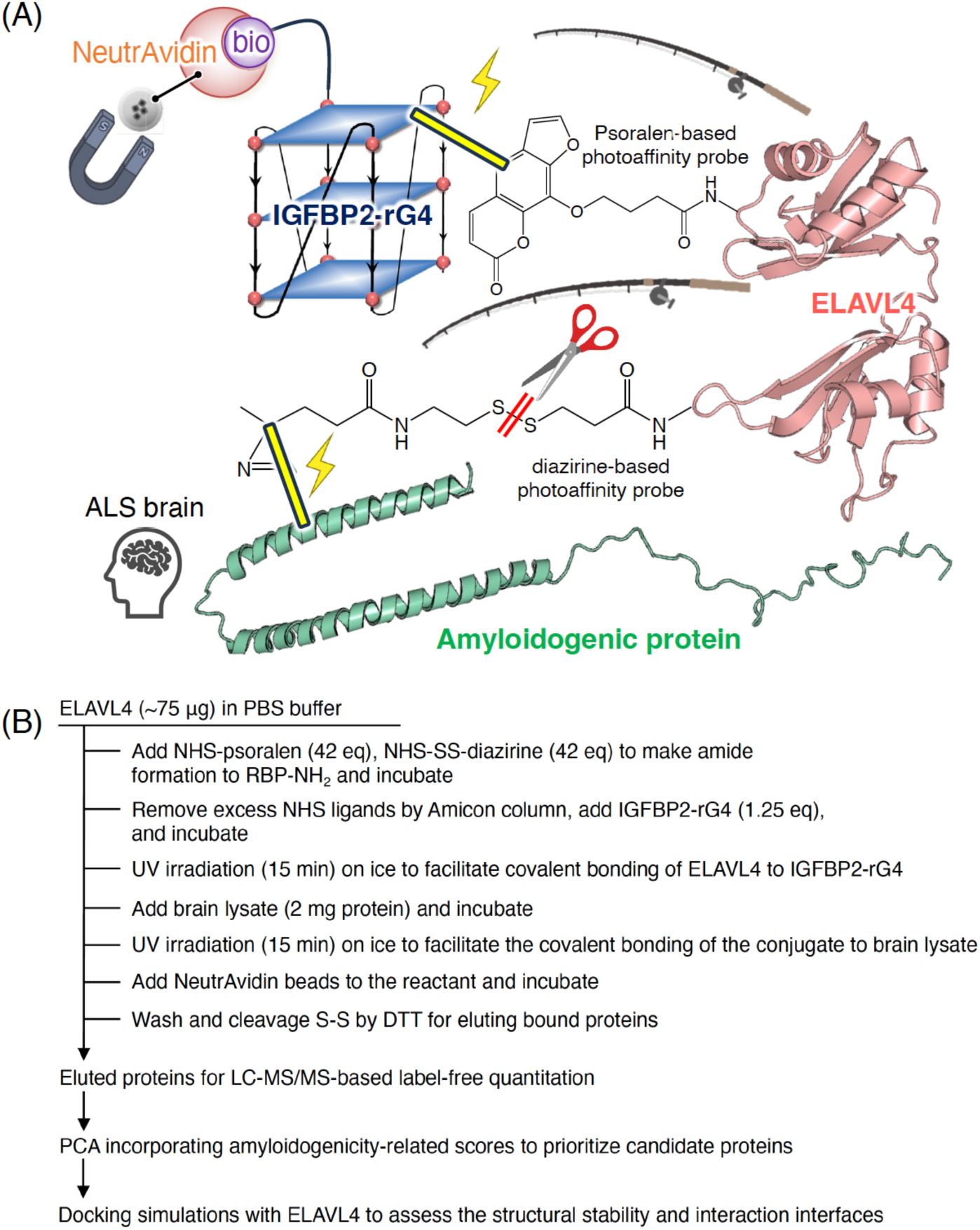
Workflow for the identification and characterization of tripartite MetaAggregates in ALS. **(A)** A combined proteomic and transcriptomic strategy was employed to search for tripartite MetaAggregates composed of RNA, RBP, and amyloids in ALS brains. Single-cell RNA-seq data from iPSC-derived motor neurons were analyzed to identify disease-associated ELAVL4 and biotinylated rG4-containing RNA sequences (IGFBP2-rG4). A synthetic photo-crosslinkable probe mimicking the tripartite MetaAggregate was generated using psoralen and diazirine chemistry, and used to pull-down binding proteins via NeutrAvidin beads. **(B)** In the workflow, the probe was incubated with brain lysates from three ALS patients or three healthy controls to capture the associated aggregative proteins. Protein identities were subsequently determined via LC-MS/MS. PCA incorporating amyloidogenicity-related scores was conducted to prioritize candidate proteins, followed by docking simulations with ELAVL4 to assess their structural stability and interaction interfaces.

## Results

### Mapping RBPs from scRNA-seq data using human iPSC motor neurons from patients with sporadic ALS

We searched for RBPs that are specifically expressed in the neurons of patients with sporadic ALS. The iPSC motor neurons from sporadic patients with ALS are particularly valuable for studying ALS’s pathogenesis; however, identifying early, neuron-specific, reproducible gene expression changes remains a challenge. In addition, bulk transcriptomic analysis lacks the necessary resolution to capture cell-type-specific gene expression changes and cellular heterogeneity, making it less accurate relative to single-cell approaches and raising the possibility of early molecular events, which are critical for understanding ALS pathology, being masked. Consequently, we selected the scRNA-seq data on iPSC-derived motor neurons deposited by Ho et al. (Ho et al, 2021) to compare the molecular characteristics of sporadic ALS patients with those of healthy control individuals.

A total of 21 pairwise comparisons between ALS and control samples were conducted. Genes were considered significantly upregulated if they met the following criteria: a log₂ fold change (log₂FC) of >0.5 and a Bonferroni-adjusted p-value of <0.05. A gene was deemed significant if it was upregulated in at least four of the 21 comparisons. This resulted in a more robust list of upregulated genes, which was used for the functional annotation and enrichment analyses. To gain deeper insights into the molecular properties of these genes, we employed multiple annotation and prediction tools. Among the 21 samples, four RBPs (ELAVL4, ZC3H13, ENO1, and LDHA) were identified, with ELAVL4 (ELAV-like protein 4) showing the highest ALS upregulation rate (**Table 1**). As a member of the ELAV family, ELAVL4 (also known as HuD) is a neuronal RBP that regulates mRNA stability, splicing, and translation, and plays critical roles in neural development, synaptic plasticity, and axonal maintenance (Akamatsu et al, 2005). Dysregulation of RNA metabolism is a hallmark of ALS, with RBPs such as ELAVL4 being increasingly implicated in disease progression (De Santis et al, 2019; Garone et al, 2021). The cytoplasmic accumulation of ELAVL4 was detected in the temporal pole regions of the postmortem ALS brain sections examined in the current study (**Fig. 3A**). These findings confirm that ELAVL4 is expressed in the pathological regions of patients.

**Figure 3.**
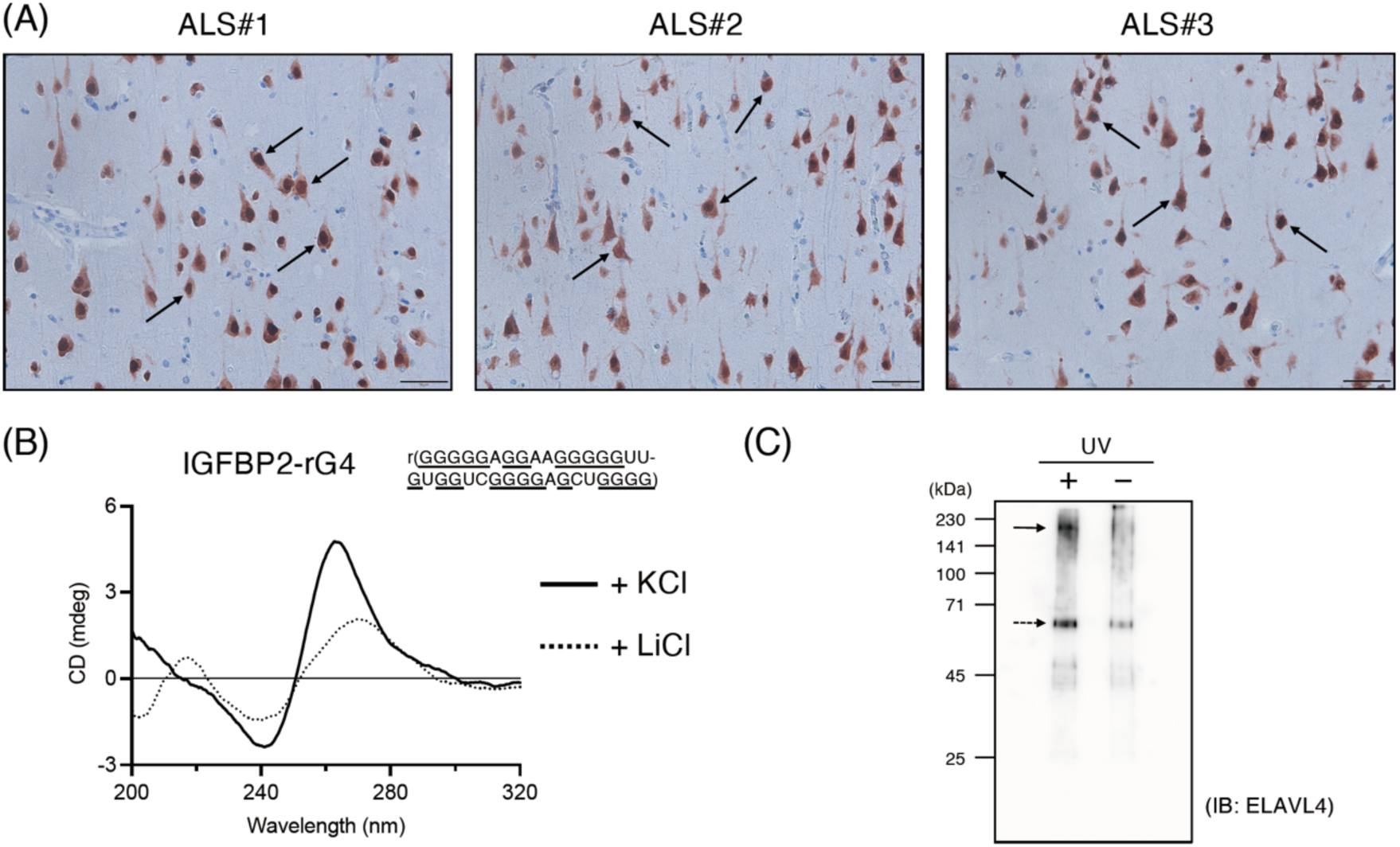
Validation of RBP and rG4 structures implicated in ALS pathology and synthesis of a MetaAggregate probe. **(A)** The brain sections from the temporal tip were stained with an anti-ELAVL4 antibody to assess its neuronal localization. Cytoplasmic accumulation (arrows) of ELAVL4 was detected in all ALS brain sections. **(B)** CD spectra of IGFBP2-rG4 (5 μM) in the presence of 100 mM KCl (solid line) or LiCl (dotted line). The guanine-rich sequence of IGFBP2-rG4 is shown. A characteristic G4 spectrum was prominently observed in the presence of the G4-promoting agent KCl, but not with the control LiCl. **(C)** Western blot of the covalent complex between ELAVL4 and IGFBP2-rG4 via psoralen/diazirine-mediated photo-crosslinking. Complexes (solid arrow) as well as unreacted ELAVL4 (dashed arrow) were immunoblotted in the photo reaction mixture (UV: +) using an anti-ELAVL4 antibody.

**Table 1.** Upregulated rG4-binding protein genes in ALS iPSC-derived motor neurons.

| Ensembl ID | Gene Symbol | Biotype | UniProt ID | MobiDB-lite<br>disorder<br>prediction | MobiDB-AlphaFold-<br>disorder | EuRBPDB<br>RBP | QUADRTlas<br>RG4<br>prediction | QUADRTlas<br>RG4<br>existence by<br>experiment | QUADRTlas<br>RG4BP<br>binding | # of samples<br>upregulated |
| --- | --- | --- | --- | --- | --- | --- | --- | --- | --- | --- |
| ENSG00000053438 | NNAT | protein_coding | Q16517 | 0.815 | - | - | TRUE | - | - | 12 |
| ENSG00000106153 | CHCHD2 | protein_coding | Q9Y6H1 | 0.735 | 0.556 | - | TRUE | - | TRUE | 5 |
| ENSG00000162374 | ELAVL4<sup>a</sup> | protein_coding | P26378 | 0.351 | 0.096 | TRUE | TRUE | - | - | 5 |
| ENSG00000148200 | NR6A1 | protein_coding | Q15406 | 0.287 | 0.188 | - | TRUE | - | TRUE | 5 |
| ENSG00000079102 | RUNX1T1 | protein_coding | Q06455 | 0.611 | 0.369 | - | TRUE | TRUE | - | 5 |
| ENSG00000115461 | IGFBP5 | protein_coding | P24593 | 0.471 | 0.074 | - | TRUE | TRUE | - | 5 |
| ENSG00000188042 | ARL4C | protein_coding | P56559 | 0.042 | - | TRUE | TRUE | TRUE | - | 4 |
| ENSG00000118432 | CNR1 | protein_coding | P21554 | 0.345 | - | - | TRUE | - | - | 4 |
| ENSG00000123200 | ZC3H13 | protein_coding | Q5T200 | 0.886 | 0.842 | TRUE | TRUE | TRUE | TRUE | 4 |
| ENSG00000251562 | MALAT1 | lncRNA | - | - | - | - | - | TRUE | TRUE | 4 |
| ENSG00000122042 | UBL3 | protein_coding | O95164 | 0.222 | - | - | TRUE | - | TRUE | 4 |
| ENSG00000109472 | CPE | protein_coding | P16870 | 0.122 | - | - | TRUE | - | TRUE | 4 |
| ENSG00000115457 | IGFBP2 | protein_coding | P18065 | 0.514 | 0.089 | - | TRUE | TRUE | TRUE | 4 |
| ENSG00000165462 | PHOX2A | protein_coding | O14813 | 0.813 | 0.493 | - | TRUE | - | - | 4 |
| ENSG00000169136 | ATF5 | protein_coding | Q9Y2D1 | 0.819 | 0.365 | - | TRUE | TRUE | TRUE | 4 |
| ENSG00000074800 | ENO1 | protein_coding | P06733 | 0 | - | TRUE | TRUE | TRUE | TRUE | 4 |
| ENSG00000134333 | LDHA | protein_coding | P00338 | 0.036 | - | TRUE | TRUE | - | TRUE | 4 |
| ENSG00000106565 | TMEM176B | protein_coding | Q3YBM2 | 0.43 | 0.126 | - | TRUE | - | - | 4 |
| ENSG00000184185 | KCNJ12 | protein_coding | Q14500 | 0.266 | 0.109 | - | TRUE | TRUE | - | 4 |
| ENSG00000133639 | BTG1 | protein_coding | P62324 | 0.275 | - | - | TRUE | TRUE | TRUE | 4 |
| ENSG00000002933 | TMEM176A | protein_coding | Q96HP8 | 0.353 | - | - | TRUE | - | - | 4 |
| ENSG00000185664 | PMEL | protein_coding | P40967 | 0.422 | 0.079 | - | TRUE | TRUE | - | 4 |

<sup>a</sup>Protein used for MetaAggregate identification.

### Mapping of the RNA sequences forming rG4 structures from scRNA-seq data using human iPSC motor neurons of sporadic ALS patients

Fragile X-associated tremor/ataxia syndrome is a neurodegenerative disorder caused by CGG-repeat expansions in the *FMR1* gene, where toxic RNA gain-of-function and RNA translation products contribute to neuronal dysfunction, in part by interacting with rG4 and potentially disrupting the normal regulatory functions of FMRP (Asamitsu et al, 2021). Based on these observations, we hypothesized that ELAVL4 contributes to ALS pathogenesis by binding to and stabilizing rG4, thereby influencing disease-relevant molecular pathways. However, the study by Ho et al. did not investigate omics-level data regarding the prevalence or expression frequency of rG4 (Ho et al., 2021).

To explore candidate rG4 sequences that may interact with ELAVL4, we employed QUADRatlas to predict rG4 structures and their putative interactions with RBPs using the scRNA-seq data on iPSC-derived motor neurons previously deposited by Ho et al. We also examined experimentally determined RNA-binding sites to validate potential RBP interactions and searched for the presence of rG4 structures using multiple G4 structure prediction models. The top 30 scoring potential G-quadruplex-forming sequences (PQSs) were selected as candidates for use in the subsequent analyses (**Table 2**). Meanwhile, considering its lack of involvement in RBPs, a 35-nt RNA sequence from IGFBP2 was identified as a possible counterpart to ELAV4 with the highest prediction score.

**Table 2.**
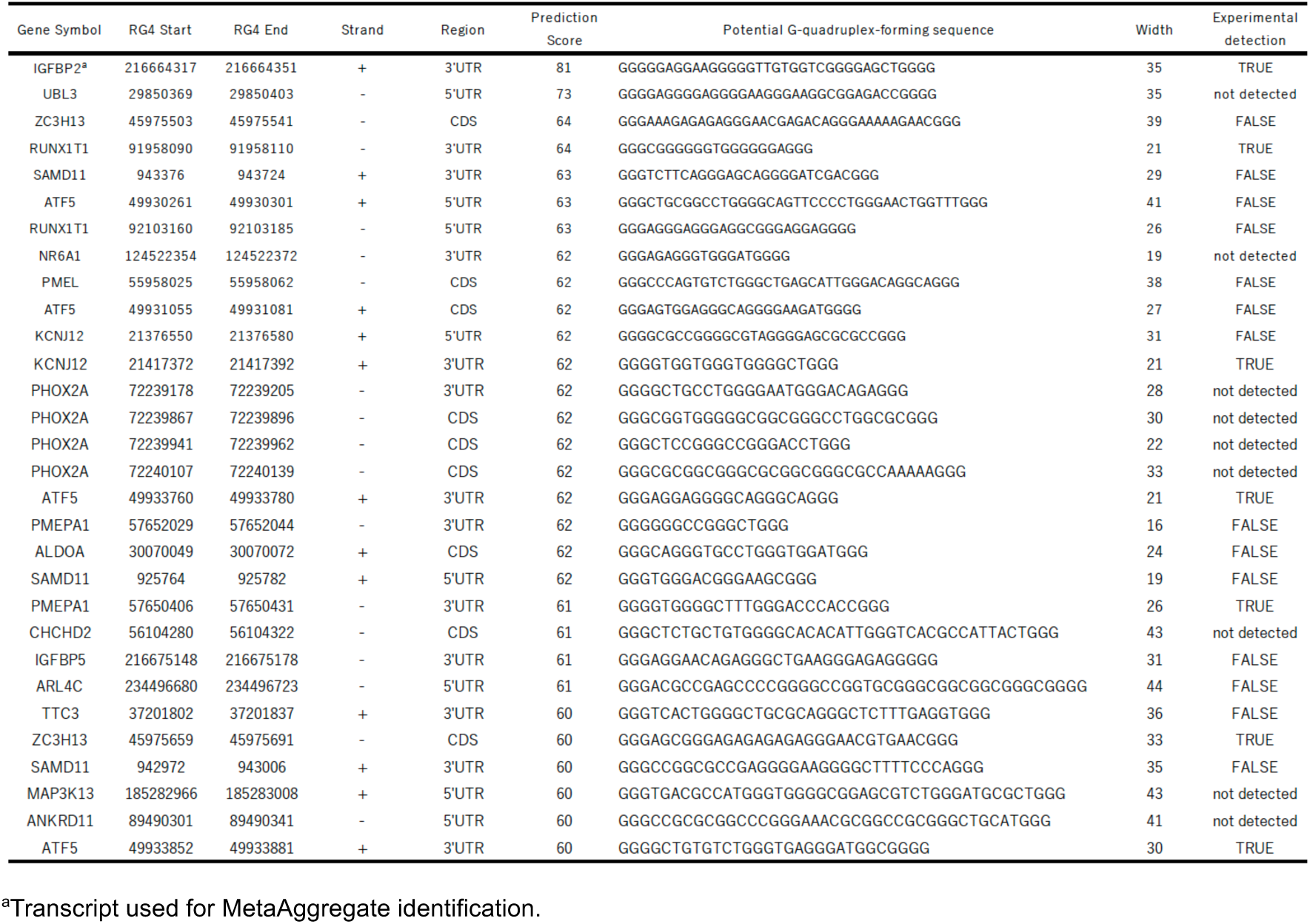
Top 30 predicted G-quadruplex-forming sequences in upregulated ALS-associated transcripts.

CD spectroscopy was carried out to confirm the structural integrity of IGFBP2 rG4. The results showed that a 35-nt RNA sequence from IGFBP2 exhibited a typical parallel-stranded rG4 in the presence of KCl, demonstrating positive and negative peaks at 262 and 240 nm, respectively, which have previously been reported to be weakly observed under LiCl conditions (**Fig. 3B**). These results indicate that a 35-nt RNA sequence from IGFBP2 can primarily fold rG4 (Del Villar-Guerra et al, 2018). This finding is consistent with the CD-NuSS data, a prediction web server of nucleic acid secondary structures from CD spectral data that uses machine learning algorithms and was previously utilized to analyze the secondary structural composition of protein (Sathyaseelan et al, 2021). The RNA mixture with random sequences (control) showed a characteristic spectrum for the stem-loop, exhibiting two positive peaks at 260 and 220 nm and a negative peak at 210 nm in the presence of KCl; this spectrum did not change even in the presence of LiCl (**Fig. 3B**). These results suggest that the 35-nt RNA sequence from IGFBP2 would form rG4 (IGFBP2-rG4).

### Search for amyloidogenic proteins composed of MetaAggregates in ALS brains

To probe the remaining amyloid of the tripartite MetaAggregate (**Fig. 1B**), we devised a pull-down strategy to link ELAVL4 to biotinylated IGFBP2-rG4. After modifying HuD-NH₂ through amide bond formation using psoralen succinimide and a diazirine succinimide containing a disulfide linkage, photo-crosslinking was employed to induce covalent bonding between ELAVL4, which was functionalized with both psoralen and diazirine groups, and IGFBP2-rG4. This process generated a synthetic MetaAggregate probe for downstream analyses (**Fig. 2A**). The complex’s formation was confirmed by Western blot (**Fig. 3C**). In this workflow, the MetaAggregate probe was first incubated with brain lysates prepared from three patients with ALS and three healthy controls. Following this incubation, a second photo-crosslinking step was carried out to covalently attach the probe to proteins in close proximity within the extracts. Finally, the disulfide bonds in the probe were cleaved under reducing conditions to release the crosslinked protein complexes for subsequent analysis (**Fig. 2B**).

LC-MS/MS-based label-free quantitation of the eluted proteins identified 79 differentially expressed proteins in the ALS brain samples, but not the healthy controls. Principal component analysis (PCA) was performed on these 79 proteins based on the scores derived from four statistical mechanics algorithms for amyloidogenicity (AmyPred-FRL, DisoRDPbind, TANGO) and intrinsic disorder (MobiDB) (**Table 3**). In the PCA plot, proteins located closer to the respective loading spots typically have higher principal component scores that reflect greater amyloidogenic potential. Accordingly, six proteins (SRP9, NDUFA2, NUFIP2, AVEN, MAL2, and SLC1A3) were selected as representative examples, as they were positioned near the loaded groups corresponding to AmyPred-FRL (#6, NDUFA2), DisoRDPbind (#1, SRP9), TANGO (#38, MAL2; #52, SLC1A3), and MobiDB (#11, NUFIP2; #34, AVEN) (**Fig. 4**).

**Figure 4.**
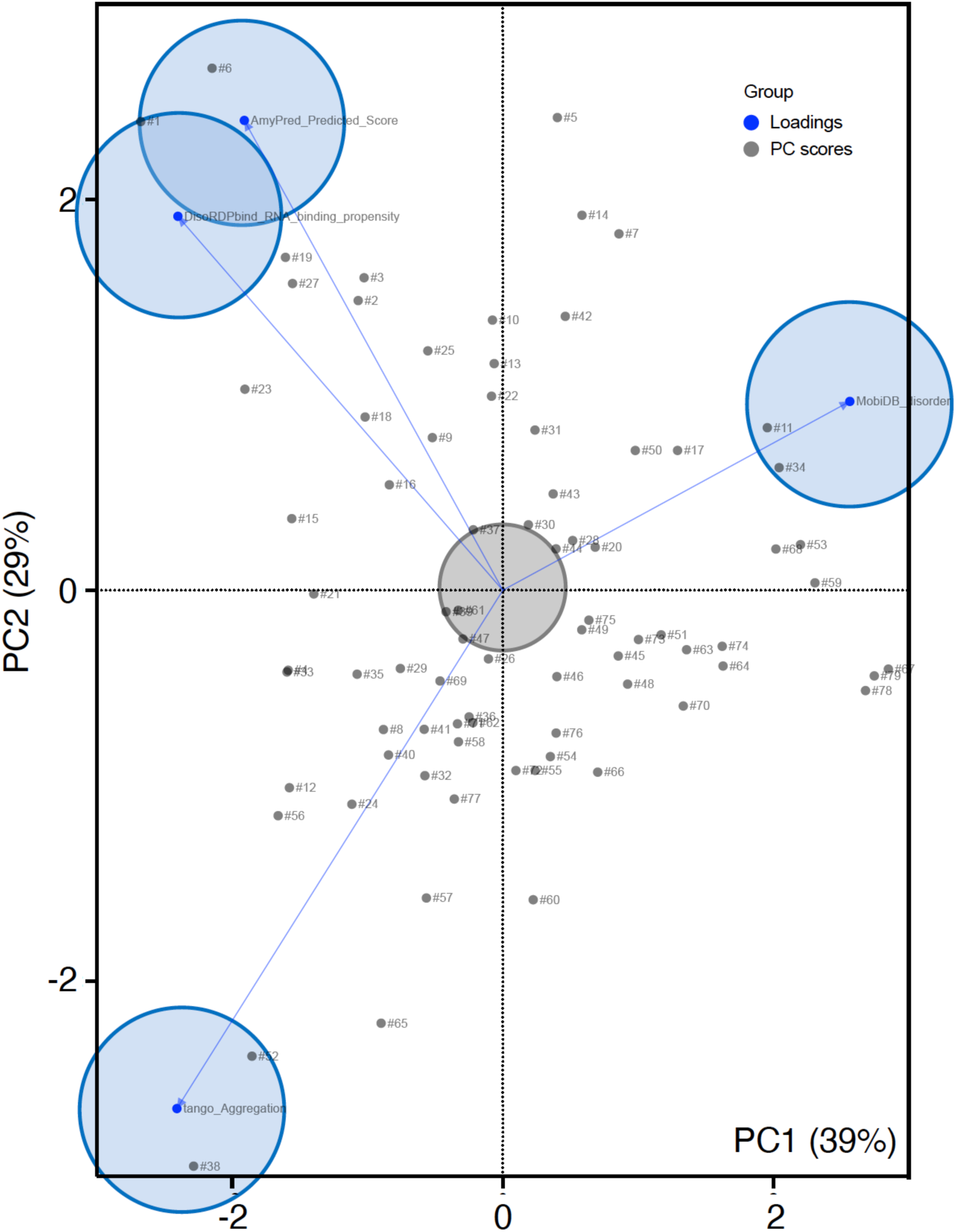
PCA of 79 candidate proteins potentially responsible for ALS pathology. PCA was performed based on amyloidogenicity and intrinsic disorder scores predicted by four independent algorithms: AmyPred-FRL, DisoRDPbind, TANGO, and MobiDB. The following six representative proteins were selected as exhibiting strong characteristic tendencies of each algorithmic cluster. These proteins were located near the corresponding loaded groups (blue circle) on the PCA plot: NDUFA2 (#6) (AmyPred-FRL), SRP9 (#1) (DisoRDPbind), MAL2 (#38) and SLC1A3 (#52) (TANGO), and NUFIP2 (#11) and AVEN (#34) (MobiDB). Proteins located near the center of the PCA plot and away from the four algorithmic clusters (gray circle) were considered to have low amyloidogenic potential and classified as non-amyloidogenic: CAPS (#37), AK3 (#39), PFDN2 (#44), STIP1 (#47), and HTRA2 (#61).

**Table 3.**
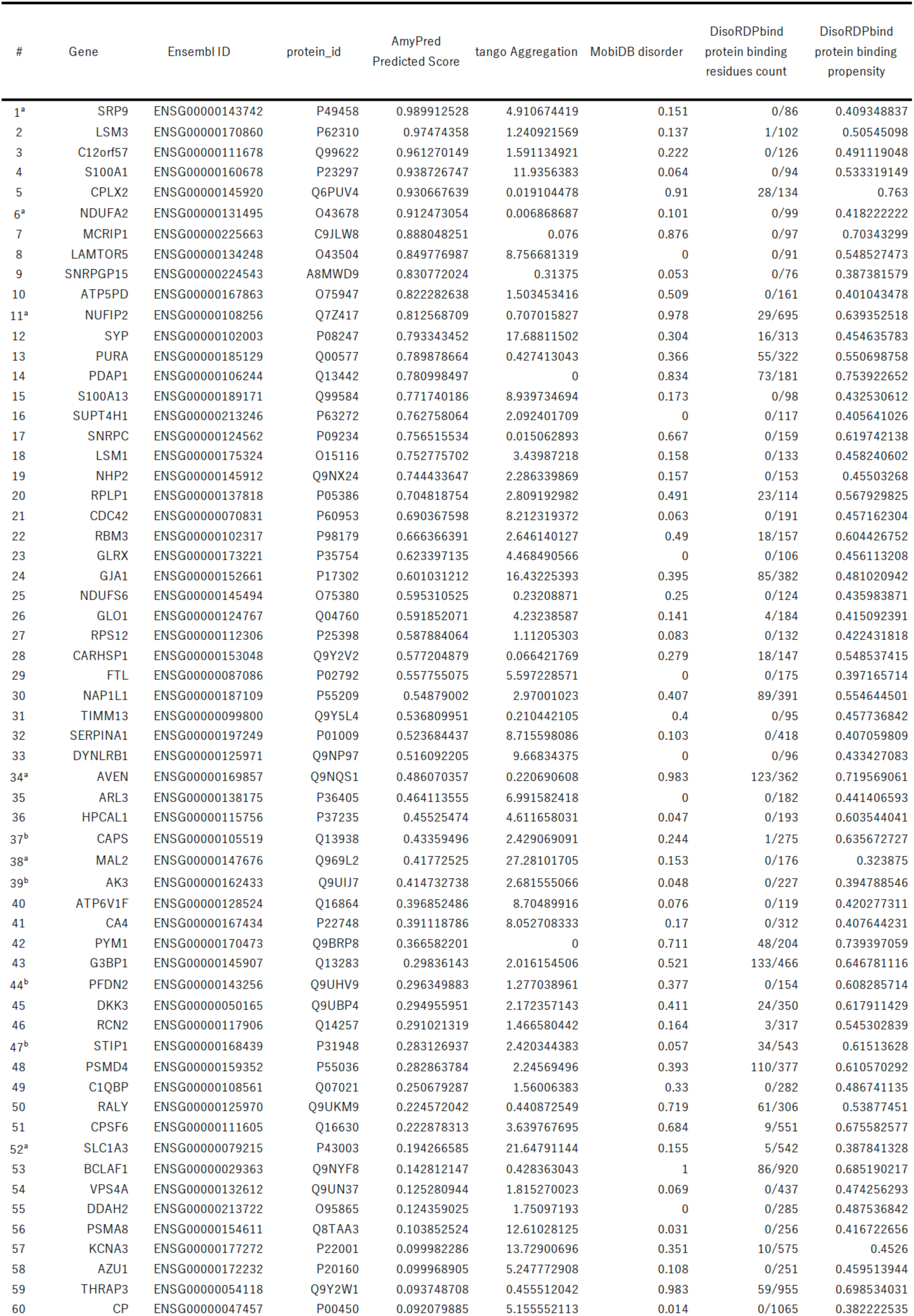
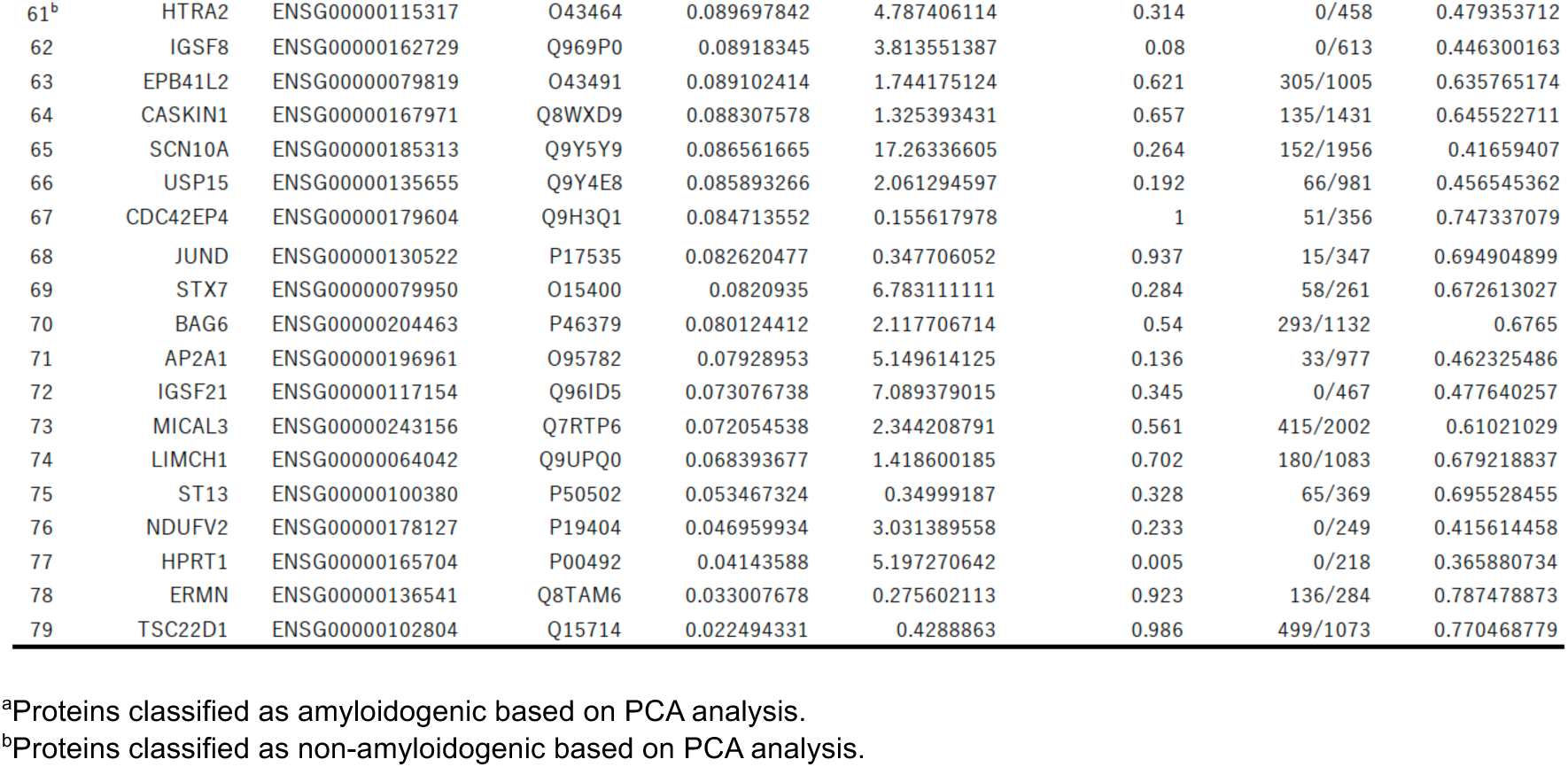
Proteins differentially expressed in ALS brains with predicted amyloidogenicity and disorder scores.

### *In silico* analysis of the structural association of ELAVL4 with amyloidogenic proteins

The potential involvement of rG4, ELAVL4, and amyloidogenic binding proteins in ALS pathology suggests that these components may coassemble into a tripartite meta-aggregate. To explore this interaction landscape and gain structural insight into the stability of these proteins, we conducted HADDOCK-based *in silico* docking to assess the structural associations between ELAVL4 and the six candidate proteins (SRP9, NDUFA2, NUFIP2, AVEN, MAL2, and SLC1A3) (**Fig. 5**). TDP-43 was used as a positive control (representing an amyloidogenic reference), while five proteins that were positioned further from the amyloid-related algorithm clusters in the PCA analysis were selected as negative controls (representing non-amyloidogenic references). All six candidate proteins exhibited strong binding to ELAVL4, with favorable HADDOCK scores (average: −95.1) and extensive interfacial contacts, particularly those involving RNA-binding motifs and intrinsically disordered regions (**Fig. 6, Table S1**). These binding affinities were similar to those observed with TDP-43 (HADDOCK score: −96.5). In contrast, negative control proteins (PFDN2, STIP1, CAPS, AK3, and HTRA2) showed much weaker interactions (average HADDOCK score: −69.2), indicating a low likelihood of specific structural association with ELAVL4. Together, these results suggest that ELAVL4 preferentially associates with proteins involved in RNA metabolism and stress granule dynamics, highlighting a potential mechanism for the formation of pathogenic aggregates in ALS.

**Figure 5.**
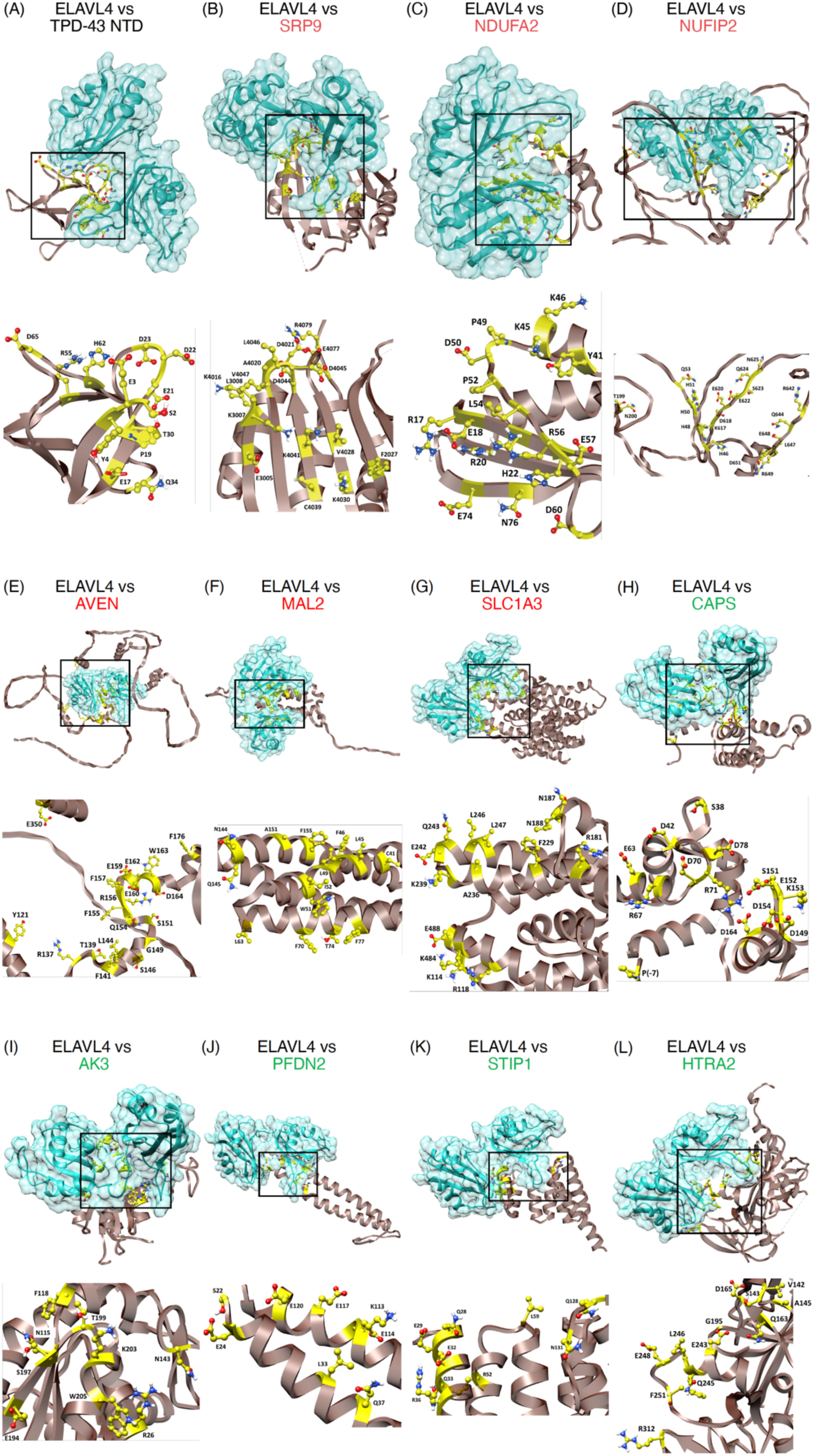
Molecular docking analysis of ELAVL4 with ALS-associated proteins. HADDOCK-based docking simulations were conducted to evaluate the structural associations between ELAVL4 (blue, PDB ID: 1FXL) and amyloidogenic proteins [red: SRP9 (PDB ID: 1914), NDUFA2 (PDB ID: 1S3A), NUFIP2 (PDB ID: AF-Q7Z417-F1), AVEN (PDB ID: AF-Q9NQS1-F1), MAL2 (PDB ID: AF-Q969L2-F1), and SLC1A3 (PDB ID: 5LLU)], which were selected based on their proximity to amyloidogenic algorithmic clusters in the PCA analysis. TDP-43 (PDB ID: 6T4B) was included as a positive control, and the five proteins with the lowest predicted amyloidogenicities [green: CAPS (PDB ID: 3E3R), AK3 (PDB ID: 6ZJB), STIP1 (PDB ID: 3Q47), HTRA2 (PDB ID: 1LCY), and PFDN2 (PDB ID: 6NR8)] were used as non-amyloidogenic proteins. The upper structure represents the most stable complex with the lowest HADDOCK score, while the lower structure provides an enlarged view of the molecular contact region (highlighted in yellow), indicated by the black rectangle.

**Figure 6.**
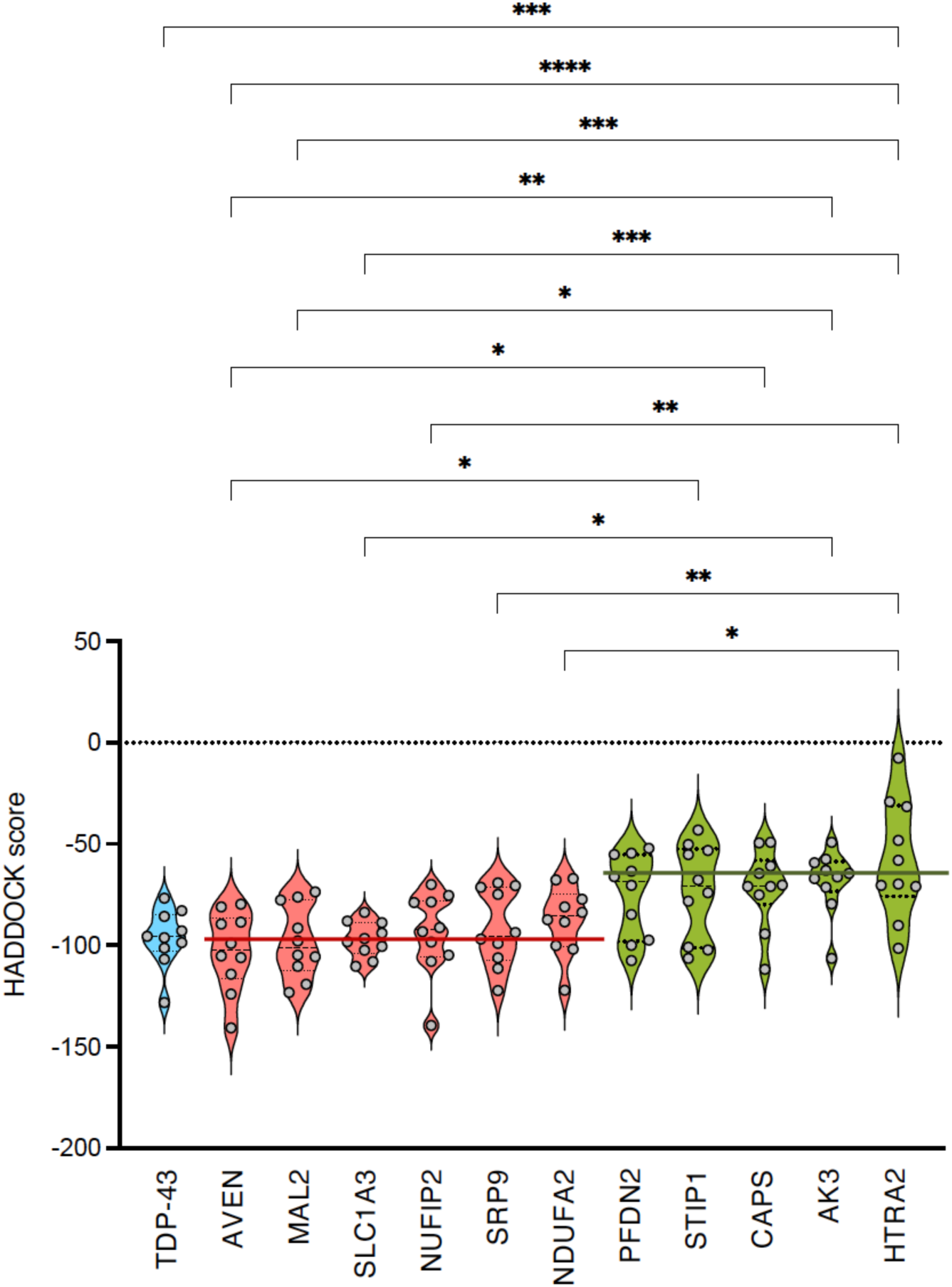
Comparison of ELAVL4 interaction interfaces with amyloidogenic (red) and non-amyloidogenic (green) proteins. All six ALS-associated proteins formed a stable complex with ELAVL4, as indicated by favorable HADDOCK scores (average: – 95.1; red solid line), similar to that of TDP-43 (–96.5). In contrast, the negative control proteins exhibited markedly weaker binding (average HADDOCK score: –69.2; green solid line), indicating reduced structural associations. *p < 0.05, **p < 0.01, ***p < 0.001, ****p < 0.0001 one-way ANOVA with post-hoc Tukey test.

## Discussion

Recent advances in scRNA-seq have provided unprecedented resolution into the cellular and molecular heterogeneity of ALS, surpassing insights offered by conventional bulk-cell transcriptomics and proteomics (Ho et al., 2021). In this study, we integrated scRNA-seq, bioinformatics, and proteomics to characterize a previously underappreciated class of heterogeneous aggregates known as MetaAggregates, which comprise RBPs, rG4s, and amyloidogenic proteins. A key finding of this study is the identification of ELAVL4, an RBP with known roles in neuronal RNA stabilization, as a core component of cytoplasmic aggregates in ALS neurons. Upregulation of ELAVL4 has been associated with neuronal dysfunction and degeneration, in part through its direct interaction with ALS-associated RNAs (De Santis et al., 2019). Later studies further demonstrated that ALS-linked FUS mutations disrupt axon growth in motor neurons and modulate the activity of ELAVL4 and Fragile X mental retardation protein (FMRP), an RBP that regulates mRNA translation, transport, and stability in neurons (Garone et al., 2021). More recently, ELAVL4 has been shown to impair neuromuscular junction integrity and induce apoptosis in human iPSC-derived neurons and *Drosophila* ALS models (Silvestri et al, 2024). The upregulation and mislocalization of ELAVL4 in ALS may reflect a compensatory response to neurodegeneration and a pathological shift in RNA metabolism. In parallel, we identified an ALS-associated rG4 sequence within the IGFBP2 transcript and utilized in a psoralen-based, biotinylated probe to search for MetaAggregates. These observations support a model in which rG4s act as scaffolds facilitating the coaggregation of RBPs and amyloidogenic proteins.

For subsequent analyses, the 30 highest-scoring PQSs were shortlisted. Additionally, an RNA sequence from IGFBP2, which readily forms an rG4 structure, was identified as the top-scoring potential partner for ELAVL4. In view of the relevance of rG4 to amyloidogenicity, several studies have previously reported G4 binding to ALS-related amyloids. The RGG domain, an arginine–glycine–glycine–rich motif, found in FUS was found to extensively recognize rG4 from r(UUAGGG)_4_, with a binding constant of *K*_D_ = 6.2 nM (Yagi et al, 2018). In subsequent studies, the RGG domain in FUS was also reported to bind rG4 deduced from post-synaptic density protein 95 (PSD-95), evaluated via steady-state fluorescence spectroscopy (*K*_D_ = 28 nM) (Imperatore et al, 2020) and surface plasmon resonance (*K*_D_ = 3.2 nM) (Ishiguro et al, 2021), respectively. Aside from rG4, other RNA structures include stem-loop RNA structures as non-canonical secondary structures, among which hnRNPA2/B1 was observed to recognize the RGG domain in FUS based on ITC analysis (*K*_D_ = 9.2 μM) (Loughlin et al, 2019). In particular, Ishiguro et al. demonstrated that liquid–liquid phase separation (LLPS) of FUS condensate formations and subsequent liquid-to-solid transition (LST) would lead to the formation of FUS aggregates through FUS’s interaction with rG4. The regulation of G4-RNA-dependent LLPS and LST pathways was lost in all ALS-linked FUS mutants tested, which were defective in rG4 recognition; this finding highlights the essential role of rG4 in these processes. Taken together, these findings suggest a mechanistic link between impaired rG4 recognition and the disruption of RBP function in the complex pathogenic cascade of ALS, and that such interactions may ultimately lead to the fibrillization of FUS.

Recent evidence suggests that ELAVL4 may play an indirect role in the pathogenesis of AD. While ELAVL4 levels have been reported to inversely correlate with Aβ accumulation, elevated ELAVL4 expression in certain AD brain regions has been associated with increased transcription of Aβ production-related genes, including *APP* and *BACE1* (Kang et al, 2014). Moreover, ELAVL4 binds to the 3′-UTRs of *APP* and *BACE1* mRNAs, stabilizing them and increasing their expression, resulting in higher Aβ levels in ELAVL4-overexpressing mice (Kang et al., 2014). ELAVL4 could be one of the stabilizing and inducing proteins of Aβ oligomerization and fibrillization (Murakami et al., manuscript in preparation). In the context of ALS, the RNA interactome of wild-type and mutant FUS using human motor neurons demonstrated that ELAVL4 interacts with FUS and is upregulated in FUS-mutant motor neuron models (De Santis et al., 2019). ELAVL4 also interacts with mutated FUS and colocalizes with FUS in cytoplasmic aggregates and stress granules (De Santis et al., 2019). Mutant FUS disrupts the normal regulation of ELAVL4, leading to ELAVL4 upregulation in a mouse primary model of ALS (Garone et al., 2021). This association of ELAVL4 with FUS indicates that ELAVL4 could be involved in coaggregation in ALS pathology.

Label-free LC-MS/MS analysis identified 79 proteins that were enriched in ALS brains, many of which are associated with synaptic dysfunction and neuroinflammation, hallmarks of ALS pathology. Proteomic studies coupled with PCA-driven analysis uncovered novel candidate amyloidogenic proteins. In particular, the PCA-prioritized proteins SRP9, NDUFA2, NUFIP2, AVEN, MAL2, and SLC1A3 exhibited strong binding to ELAVL4, similar to that observed for TDP-43, and displayed structural features suggestive of amyloidogenic potential. Below, we summarize whether and how these six proteins are implicated in ALS;

- **SRP9**, a component of the signal recognition particle (SRP), forms a heterodimer with SRP14 to constitute the Alu domain of SRP and is involved in the cotranslational targeting of nascent peptides from the ribosome to the endoplasmic reticulum (Amemiya et al, 1977). Although the impaired cotranslation from the ribosome to the endoplasmic reticulum may be related to ALS pathology (Spencer et al, 1997), its direct involvement in ALS remains unclear.
- **NDUFA2**, a subunit of mitochondrial complex I, is involved in ATP production through the mitochondrial electron transfer chain. While mitochondrial dysfunction is central to ALS (Spencer et al., 1997), direct evidence implicating NDUFA2 in ALS pathogenesis is scant.
- **NUFIP2** an RBP identified as a potent interactor of ATXN2L in neural development, is functionally involved in RNA granule trafficking and stress granule regulation. NUFIP2, highlighted as the most miRNA-targeted gene in the integrative analysis, plays a regulatory role in ALS pathogenesis through extensive miRNA-mediated gene control (Hamzeiy et al, 2018). The contribution of NUFIP2 to ALS pathogenesis through interaction with ATXN2L and aggregation-prone proteins has been implicated previously (Key et al, 2025). NUFIP2 contributes to ALS pathology by interacting with mislocalized TDP-43, promoting its sequestration into cytoplasmic aggregates and co-localizing with TDP-43 pathology in patient tissues (Xie et al, 2025).
- **AVEN** an apoptosis and RNA splicing regulator (Chau et al, 2000), contributes in part to ALS pathology by blocking apoptosis in motor neurons, as its upregulation was associated with caspase-independent neuronal degeneration in an ALS model carrying G93A-SOD1 (Martin et al, 2007).
- **MAL2** is a protein essential for membrane trafficking and lipid raft maintenance (Li et al, 2024). Although its functional disruption and protein trafficking defects are prominent features in ALS, direct evidence linking MAL2 to the disease is currently lacking.
- **SLC1A3** is a glutamate transporter responsible for the uptake of extracellular glutamate and plays a key role in preventing excitotoxicity (Kanai & Hediger, 2004). In a G93A-SOD1-carrying ALS model, SLC1A3 was upregulated in astrocytes and functioned as a negative regulator of potassium uptake, leading to impaired K⁺ buffering capacity, elevated extracellular potassium levels, and neuronal hyperexcitability, thereby promoting neurodegeneration (Ding et al, 2024).

Gene Ontology analysis further highlighted significant enrichment in pathways related to intracellular membrane systems or involved in transport, nucleic acid binding, or developmental processes. Their functions suggest a shared role in maintaining cellular homeostasis through membrane-associated activities and gene expression regulation. Among these, NUFIP2 (Hamzeiy et al., 2018; Key et al., 2025), AVEN (Martin et al., 2007), and SLC1A3 (Ding et al., 2024) have been previously implicated in ALS pathology. It should be noted that the detection of multiple RBPs and amyloid-like proteins beyond TDP-43 indicates that ALS involves a broader spectrum of aggregation-prone proteins than previously recognized. Mechanistically, the incorporation of ELAVL4 into cytoplasmic aggregates may lead to widespread disruption of RNA processing, akin to TDP-43 or FUS misbehavior during dysregulated liquid–liquid phase separation. Importantly, our results align with a previous report demonstrating that rG4s facilitate αSyn aggregation (Matsuo et al., 2024), suggesting that rG4–RBP interactions can act as nucleation platforms for MetaAggregate formation. From a translational perspective, the rG4-based probe developed in this study offers a promising tool for the detection of MetaAggregates in patient tissues. Further work is needed to validate its utility in biofluid-based biomarkers (e.g., CSF or blood). Characterization of the pharmacological modulation of rG4–RBP interactions may provide a novel therapeutic approach, inspired by recent efforts targeting rG4s in other disease contexts.

In conclusion, this study proposes a set of previously unrecognized amyloidogenic protein candidates potentially involved in ALS pathogenesis. Identified through proteomic screening and PCA-guided analysis, these proteins exhibit strong binding affinity to ELAVL4 and possess sequence or structural features consistent with amyloid-forming propensity. Many are functionally linked to RNA metabolism or membrane trafficking, suggesting that such molecular pathways may harbor novel contributors to pathogenic protein aggregation. While their individual relevance to disease remains to be validated, this approach offers a framework for the systematic identification of pathogenic amyloids in sporadic ALS. More broadly, it may also inform investigations about mixed pathology mechanisms and contribute to biomarker discovery efforts.

## Methods

### Bioinformatics

First, we used GENCODE v30 (https://www.gencodegenes.org/) to classify genes as coding or non-coding. In addition, to assess intrinsic disordered regions and identify RBPs, protein domain analysis was performed using MobiDB (https://mobidb.org/) and EuRBPDB (http://eurbpdb.gzsys.org.cn/index.php) (Liao et al, 2020), respectively. To further refine the selection of rG4-binding proteins, we also incorporated QUADRatlas data (https://rg4db.cibio.unitn.it/) (Bourdon et al, 2023). PQSs used in the experiments were obtained by selecting the genes that are upregulated in ALS pathogenesis. The scRNA-seq data of iPSC-derived motor neurons from healthy individuals was selected along with published data obtained previously from sporadic and familial ALS patients (Ho et al., 2021). Filtering of the DEGs (differentially expressed genes) was performed by selecting those that were upregulated with a log fold change (log FC) greater than 0.5 and a Bonferroni-adjusted p-value less than 0.05. Genes that were upregulated in at least four out of the 21 sample comparisons were retained, resulting in 22 candidate genes for the downstream analysis (**Table 1**). For each of the selected genes, the EuRBPDB database was used to assess whether the corresponding protein product could be an RBP. To determine whether the proteins contained intrinsically disordered regions, they were evaluated using the Mobi database (MobiDB) (Piovesan et al, 2025).

The QUADRatlas database was used to investigate the presence of known or predicted rG4s within the transcripts of the selected genes. We also examined experimentally determined RNA-binding sites based on ENCODE eCLIP assays to validate potential RBP interactions. To search for the presence of rG4 structures, we categorized the rG4s into experimentally determined and predicted rG4s. Experimentally validated rG4s were identified using RT-stop profiling and rG4-seq techniques, which are designed to detect RNA secondary structures, while predicted rG4s were assessed using multiple computational tools with the following thresholds: QGRS Mapper, score threshold = 19; pqsfinder score threshold = 47; G4Hunter score threshold = 1.2, window size = 25). Additionally, to determine rG4-binding proteins, we utilized RNA pull-down experiments and curated evidence from the existing literature to validate their functional relevance. Together, these analyses provided a comprehensive understanding of the transcriptomic landscape in ALS, highlighting key upregulated genes, their rG4-forming propensity, and potential RNA-protein interactions that may contribute to disease pathogenesis. In addition, PQSs were identified using QGRS Mapper (Kikin et al, 2006), and the top 30 scoring PQSs were selected as candidates for use in the subsequent analyses. **Table 2** represents ALS-specific genes with a high propensity to form rG4 structures (with a prediction score of 60 or higher). Among them, IGFBP2 showed high scores across multiple G4 structure prediction models (QGRS Mapper, pqsfinder, and G4Hunter).

### Histology and immunohistochemistry

For each brain, samples from the frontal tip, temporal tip, occipital tip, and cerebellar hemisphere were frozen for the biochemical analyses. Brain samples were fixed in 10% buffered formalin, embedded in paraffin, and sliced into 6-μm-thick sections. Histological staining included the hematoxylin–eosin, Klüver–Barrera, Bodian, Gallyas–Braak, and Holzer methods. Immunohistochemical staining was performed using the primary antibody against ELAVL4 (1:100, mouse monoclonal, clone E-1; Santa Cruz, Dallas, TX, USA).

### Circular dichroism spectroscopy

CD spectra were measured using a JASCO J-805 spectroscope (JASCO, Tokyo, Japan) as described previously (Murakami et al, 2022b) with a slight modifications. For RNA measurement, the refolded RNA solution (5 μM, 200 μL), along with 10 mM Tris-HCl (pH 7.5) containing 100 mM KCl or LiCl was loaded into the 1-mm pathlength quartz cell (JASCO). The CD spectrum was recorded (scanning speed: 100 nm/min) over the wavelength range of 200–320 nm for RNA measurement. The spectra were mapped after subtraction of the spectrum of the vehicle.

### Synthesis of ELAVL4 probe

To create the ELAVL4 probe, we added a solution of His-ELAVL4 (3.2 nmole, abcam, Cambridge, UK) in D-PBS(−) (Nacalai, Japan) to a solution of NHS-psoralen (16 nmole; Thermo, Waltham, MA, USA) and sulfo-SDAD (16 nmole, Thermo) in D-PBS(-) in 1.5 mL Protein LoBind^®^ tubes. After incubation in a rotator shielded from light for 1 h at 25°C, the unreacted ligands were removed via centrifugation with a 3-kDa cutoff value (Merck Millipore, Burlington, MA, USA). A biotinylated 35-nt RNA sequence from IGFBP2 in D-PBS(−) (GGGGGAGGAAGGGGGUU-GUGGUCGGGGAGCUGGGG; IGFBP2-rG4, FASMAC, Kanagawa, Japan) was added to the fraction containing His-ELAVL4. After incubation with the same conditions stated above, the resulting mixture was UV-irradiated (365 nm, 100W; Thermo) on ice for 15 min to conjugate ELAVL4 with IGFBP2-rG4. The purity of the solution was then validated via SDS-PAGE.

### Western blotting

Crosslinked ELAVL4 (2.5 μg) were treated with 4× LDS sample buffer (Invitrogen; Carlsbad, CA, USA) before heating at 70°C for 10 min. The denatured samples were separated via SDS-PAGE on NuPAGE 10% Bis-Tris gel (Invitrogen) and subsequent transfer to PVDF membranes (0.45 μm pore size, Merck Millipore). The membranes were blocked and incubated with the primary antibody of anti-ELAVL4 (E-1, Santa Cruz; 0.2 μg/mL). Following primary antibody incubation, blots were washed before being incubated with the appropriate secondary antibody. Blots were developed with enhanced chemiluminescence and imaged with Lumino Graph II (ATTO; Tokyo, Japan).

### Patient consent and ethics declaration

Tissue samples were obtained at the National Hospital Organization, Iou National Hospital. Neuropathologically, three cases were diagnosed with ALS, and another three cases without ALS pathology were employed as controls (**Table 4**). This study was approved by the Medical Ethics Review Board of Kanazawa University and Kyoto University (approval numbers 714319 and 114487). Informed consent was obtained from the patients as an opt-out on the website. All experiments involving human material were conducted in accordance with the Declaration of Helsinki.

**Table 4.** Clinical and pathological information of the patients^a^.

| Patient | Age | Sex | Brettschneider stage | Braak amyloid stage | CERAD | Thal phase | Braak NFT stage |
| --- | --- | --- | --- | --- | --- | --- | --- |
| ALS#1 | 56 | F | 4 | 0 | None | 0 | I |
| ALS#2 | 84 | F | 4 | 0 | None | 0 | II |
| ALS#3 | 77 | M | 4 | B | Sparse | 3 | II |
| DC#1<br>(Lewy body disease) | 71 | M | N/A | B | None | 2 | II |
| DC#2<br>(Lewy body disease) | 67 | M | N/A | A | None | 2 | I |
| DC#3<br>(Myotonic dystrophy) | 64 | M | N/A | 0 | None | 0 | III-IV |

<sup>a</sup>ALS, amyotrophic lateral sclerosis; CERAD, consortium to establish a registry for Alzheimer's disease; DC, disease control; N/A, not available; NFT, neurofibrillary tangles.

### Tissue preparation

Human brain tissue (temporal pole, 0.2–0.3 g) was homogenized in 10 volumes (w/v) of PBS (pH 7.4) containing a mixture of protease and phosphatase inhibitors (Nacalai Tesque, Kyoto, Japan), 0.1 µM pepstatin A, 1 mM phenylmethylsulfonyl fluoride, 0.5% Tween-20, and 0.5% Triton X-100. The homogenates were centrifuged at 17,000 g for 30 min at 4°C to obtain a supernatant (PBS-soluble) fraction with a sufficient protein concentration for the subsequent analyses (2–3 mg/mL). The total protein concentration of the brain lysates was determined via the Bradford protein assay (Bio-Rad; Hercules, CA, USA).

### Photolabeling of ELAVL4 binding proteins in ALS brain lysates

The brain lysates (2 mg/mL, 1 mL) were added to a solution of ELAVL4 probes (3.2 nmole). After incubation in a rotator for 1 h at room temperature in the dark, the solution was added to the FG-NeutrAvidin beads (1 mg, Tamagawa Seiki, Japan) in 1.5-mL Protein LoBind^®^ tubes. The resulting mixture was UV-irradiated (365 nm, 100W; Thermo) on ice for 15 min to conjugate the complex (ELAVL4+IGFBP2-rG4) with the proteins in the brain lysates. For disulfide cleavage and elution from the beads, RIPA buffer (Nacalai, Japan) containing 50 mM dithiothreitol was incubated at 60°C with shaking at 1,200 rpm for 1 h after washing with D-PBS(−), and the supernatants were collected and subjected to mass spectrometry.

### Protein identification using nano-liquid chromatography mass spectrometry

The trypsin-digested peptides were analyzed by Orbitrap QE plus (Thermo Fisher Scientific) with nano-liquid chromatography (EASY-nLC 1200; Thermo Fisher Scientific). The purified peptides were loaded and separated on the Aurora column (25 cm x 75 µm ID, 1.6 mm C18; Ionoptics) with a linear acetonitrile gradient (0-35%) in 0.1% formic acid at a flow rate of 300 nL min-1. The peptide ions were detected by Orbitrap QE plus MS; Thermo Fisher Scientific) in the data-dependent acquisition mode with the installed Xcalibur software (Thermo Fisher Scientific). Full-scan mass spectra were acquired in the MS over 375-1,500 m/z with resolution of 70,000. The MS/MS analyses were carried out using SEQUEST HT search algorithms against the Homo sapiens (Swiss prot. Tax ID 9609) protein database (2024-11-30) using Proteome Discoverer (PD) 3.0 (Version 3.0.1.27; Thermo Fisher Scientific; Waltham, MA, USA). Label-free quantification was also performed with PD 3.0 using precursor ions quantifier nodes. The processing workflow included spectrum files RC, spectrum selector, SEQUEST HT search nodes, percolator, ptmRS, and minor feature detector nodes. Oxidation of methionine were set as a variable modification and carbamidomethylation of cysteine was set as a fixed modification. Mass tolerances in MS and MS/ MS were set at 10 ppm and 0.6 Da, respectively. Trypsin was specified as protease and a maximum of two missed cleavages were allowed. Target-decoy database searches used for calculation of false discovery rate (FDR) and for peptide identification FDR was set at 1%. Label-free quantification was also performed with PD 3.0 using precursor ions quantifier nodes. The consensus workflow included MSF files, Feature Mapper, precursor ion quantifier, PSM groper, peptide validator, peptide and protein filter, protein scorer, protein marker, protein FDR validator, protein grouping, peptide in protein. Normalization of the abundances was performed using total peptide amount mode. The accession number of protein mass spectrometry data is PXD065772 for ProteomeXchange and JPST003914 for jPOST. Identified proteins are listed in **Table 3**.

### *In silico* molecular docking

We evaluated the association of ELAVL4 (PDB code: 1FXL) with the following proteins, whose PDB entries were selected with a long solved-structure region: N-terminal domain of TDP-43 (PDB: 6T4B), SRP9 (#1, PDB: 1914), NDUFA2 (#6, PDB: 1S3A), NUFIP2 (#11, PDB: AF-Q7Z417-F1), AVEN (#34, PDB: AF-Q9NQS1-F1), CAPS (#37, PDB: 3E3R), MAL2 (#38, PDB: AF-Q969L2-F1), AK3 (#39, PDB: 6ZJB), PFDN2 (#44, PDB: 6NR8), STIP1 (#47, PDB: 3Q47), SLC1A3 (#52, PDB: 5LLU), and HTRA2 (#61, PDB: 1LCY). The docking calculations were performed using HADDOCK version 2.4 software (Dominguez et al, 2003; Honorato et al, 2021; Honorato et al, 2024). In order to optimize the docking program, the water, solvents, and nucleic acids were removed from the protein structures. The HADDOCK score, van der Waals energy values, Z-score, and root mean square deviation were collected to evaluate the protein-protein interactions. The top ten structures were generated for each docking process, and the HADDOCK scores were used to select the best scoring models.

### Statistical analysis

Statistical analysis was performed using the scientific data analysis software GraphPad Prism version 6 (GraphPad Software) with one-way analysis of variance (ANOVA) followed by post-hoc Tukey test. p < 0.05 represented a statically significant difference, which is indicated in each figure legend. All statical analyses were performed using the GraphPad Prism 9.3.1 (GraphPad Software, San Diego, CA).

## Acknowledgments

The authors acknowledge funding in part by JSPS KAKENHI, grant number 23H03852 to K.M. We thank Ms. Ritsuko Goto (Kanazawa University) for providing technical support and enago (https://www.enago.jp/) for English editing.

## Author contributions

Kazuma Murakami supervised, contributed to the conception and design of the study, and wrote the initial draft of the manuscript. Kazuma Murakami, Thi-Hong-Van Nguyen, Nobuko Fujita, Chioko Nagano, and Takumi Nishiuchi performed the experiments. Kenji Mizuguchi, Yasuhiro Sakashita, Moeko Noguchi-Shinohara, and Kenjiro Ono contributed to the resource. Kenjiro Ono supervised the study. The manuscript was written and edited based on the contributions of all authors. All authors have read and approved the final version of the manuscript.

## Data Availability

The accession number of protein mass spectrometry data is PXD065772 for ProteomeXchange and JPST003914 for jPOST. The data that support the findings of this study are available from the corresponding author (K.M.) upon reasonable request. Supplementary information is available for this paper at https://doi.org/XXX.

## REFERENCES

Akamatsu W, Fujihara H, Mitsuhashi T, Yano M, Shibata S, Hayakawa Y, Okano HJ, Sakakibara S, Takano H, Takano T et al (2005) The RNA-binding protein HuD regulates neuronal cell identity and maturation. Proc Natl Acad Sci U S A 102: 4625–4630

Amador-Ortiz C, Lin WL, Ahmed Z, Personett D, Davies P, Duara R, Graff-Radford NR, Hutton ML, Dickson DW (2007) TDP-43 immunoreactivity in hippocampal sclerosis and Alzheimer’s disease. Ann Neurol 61: 435–445

Amemiya E, Soma A, Iida S (1977) [Outlook and trends in reports presented to the Adult Nursing Section of the Japan Nursing Academic Society in the past 3 years - 1972, 1973, and 1974]. Kango Kenkyu 10: 143-145

Apicco DJ, Ash PEA, Maziuk B, LeBlang C, Medalla M, Al Abdullatif A, Ferragud A, Botelho E, Ballance HI, Dhawan U et al (2018) Reducing the RNA binding protein TIA1 protects against tau-mediated neurodegeneration in vivo. Nat Neurosci 21: 72–80

Arai T, Hasegawa M, Akiyama H, Ikeda K, Nonaka T, Mori H, Mann D, Tsuchiya K, Yoshida M, Hashizume Y et al (2006) TDP-43 is a component of ubiquitin-positive tau-negative inclusions in frontotemporal lobar degeneration and amyotrophic lateral sclerosis. Biochem Biophys Res Commun 351: 602–611

Asamitsu S, Yabuki Y, Ikenoshita S, Kawakubo K, Kawasaki M, Usuki S, Nakayama Y, Adachi K, Kugoh H, Ishii K et al (2021) CGG repeat RNA G-quadruplexes interact with FMRpolyG to cause neuronal dysfunction in fragile X-related tremor/ataxia syndrome. Sci Adv 7

Balendra R, Isaacs AM (2018) C9orf72-mediated ALS and FTD: multiple pathways to disease. Nat Rev Neurol 14: 544–558

Bayer TA (2015) Proteinopathies, a core concept for understanding and ultimately treating degenerative disorders? Eur Neuropsychopharmacol 25: 713–724

Benilova I, Karran E, De Strooper B (2012) The toxic Aβ oligomer and Alzheimer’s disease: an emperor in need of clothes. Nat Neurosci 15: 349–357

Bitan G, Fradinger EA, Spring SM, Teplow DB (2005) Neurotoxic protein oligomers--what you see is not always what you get. Amyloid 12: 88–95

Bourdon S, Herviou P, Dumas L, Destefanis E, Zen A, Cammas A, Millevoi S, Dassi E (2023) QUADRatlas: the RNA G-quadruplex and RG4-binding proteins database. Nucleic Acids Res 51: D240–D247

Brettschneider J, Del Tredici K, Lee VM, Trojanowski JQ (2015) Spreading of pathology in neurodegenerative diseases: a focus on human studies. Nat Rev Neurosci 16: 109–120

Brown RH, Al-Chalabi A (2017) Amyotrophic Lateral Sclerosis. N Engl J Med 377: 162–172

Chau BN, Cheng EH, Kerr DA, Hardwick JM (2000) Aven, a novel inhibitor of caspase activation, binds Bcl-xL and Apaf-1. Mol Cell 6: 31–40

Colom-Cadena M, Gelpi E, Charif S, Belbin O, Blesa R, Marti MJ, Clarimon J, Lleo A (2013) Confluence of alpha-synuclein, tau, and beta-amyloid pathologies in dementia with Lewy bodies. J Neuropathol Exp Neurol 72: 1203–1212

De Santis R, Alfano V, de Turris V, Colantoni A, Santini L, Garone MG, Antonacci G, Peruzzi G, Sudria-Lopez E, Wyler E et al (2019) Mutant FUS and ELAVL4 (HuD) Aberrant Crosstalk in Amyotrophic Lateral Sclerosis. Cell Rep 27: 3818–3831 e3815

Del Villar-Guerra R, Trent JO, Chaires JB (2018) G-Quadruplex Secondary Structure Obtained from Circular Dichroism Spectroscopy. Angew Chem Int Ed Engl 57: 7171–7175

Ding F, Sun Q, Long C, Rasmussen RN, Peng S, Xu Q, Kang N, Song W, Weikop P, Goldman SA et al (2024) Dysregulation of extracellular potassium distinguishes healthy ageing from neurodegeneration. Brain 147: 1726–1739

Dominguez C, Boelens R, Bonvin AM (2003) HADDOCK: a protein-protein docking approach based on biochemical or biophysical information. J Am Chem Soc 125: 1731–1737

Garone MG, Birsa N, Rosito M, Salaris F, Mochi M, de Turris V, Nair RR, Cunningham TJ, Fisher EMC, Morlando M et al (2021) ALS-related FUS mutations alter axon growth in motoneurons and affect HuD/ELAVL4 and FMRP activity. Commun Biol 4: 1025

Gilpin KM, Chang L, Monteiro MJ (2015) ALS-linked mutations in ubiquilin-2 or hnRNPA1 reduce interaction between ubiquilin-2 and hnRNPA1. Hum Mol Genet 24: 2565–2577

Goedert M (2020) Tau proteinopathies and the prion concept. Prog Mol Biol Transl Sci 175: 239–259

Haeusler AR, Donnelly CJ, Rothstein JD (2016) The expanding biology of the C9orf72 nucleotide repeat expansion in neurodegenerative disease. Nat Rev Neurosci 17: 383–395

Hamzeiy H, Suluyayla R, Brinkrolf C, Janowski SJ, Hofestadt R, Allmer J (2018) Visualization and Analysis of miRNAs Implicated in Amyotrophic Lateral Sclerosis Within Gene Regulatory Pathways. Stud Health Technol Inform 253: 183–187

Hasegawa K, Yamaguchi I, Omata S, Gejyo F, Naiki H (1999) Interaction between Aβ(1-42) and Aβ(1-40) in Alzheimer’s β-amyloid fibril formation in vitro. Biochemistry 38: 15514–15521

Hellstrand E, Boland B, Walsh DM, Linse S (2010) Amyloid β-protein aggregation produces highly reproducible kinetic data and occurs by a two-phase process. ACS Chem Neurosci 1: 13–18

Henderson MX, Trojanowski JQ, Lee VM (2019) alpha-Synuclein pathology in Parkinson’s disease and related alpha-synucleinopathies. Neurosci Lett 709: 134316

Higashi S, Iseki E, Yamamoto R, Minegishi M, Hino H, Fujisawa K, Togo T, Katsuse O, Uchikado H, Furukawa Y et al (2007) Concurrence of TDP-43, tau and alpha-synuclein pathology in brains of Alzheimer’s disease and dementia with Lewy bodies. Brain Res 1184: 284–294

Ho R, Workman MJ, Mathkar P, Wu K, Kim KJ, O’Rourke JG, Kellogg M, Montel V, Banuelos MG, Arogundade OA et al (2021) Cross-Comparison of Human iPSC Motor Neuron Models of Familial and Sporadic ALS Reveals Early and Convergent Transcriptomic Disease Signatures. Cell Syst 12: 159–175 e159

Honorato RV, Koukos PI, Jimenez-Garcia B, Tsaregorodtsev A, Verlato M, Giachetti A, Rosato A, Bonvin A (2021) Structural Biology in the Clouds: The WeNMR-EOSC Ecosystem. Front Mol Biosci 8: 729513

Honorato RV, Trellet ME, Jimenez-Garcia B, Schaarschmidt JJ, Giulini M, Reys V, Koukos PI, Rodrigues J, Karaca E, van Zundert GCP et al (2024) The HADDOCK2.4 web server for integrative modeling of biomolecular complexes. Nat Protoc 19: 3219–3241

Ilie IM, Caflisch A (2019) Simulation studies of amyloidogenic polypeptides and their aggregates. Chem Rev 119: 6956–6993

Imperatore JA, McAninch DS, Valdez-Sinon AN, Bassell GJ, Mihailescu MR (2020) FUS Recognizes G Quadruplex Structures Within Neuronal mRNAs. Front Mol Biosci 7: 6

Irwin DJ, Lee VM, Trojanowski JQ (2013) Parkinson’s disease dementia: convergence of alpha-synuclein, tau and amyloid-beta pathologies. Nat Rev Neurosci 14: 626–636

Ishiguro A, Lu J, Ozawa D, Nagai Y, Ishihama A (2021) ALS-linked FUS mutations dysregulate G-quadruplex-dependent liquid-liquid phase separation and liquid-to-solid transition. J Biol Chem 297: 101284

Ito D, Hatano M, Suzuki N (2017) RNA binding proteins and the pathological cascade in ALS/FTD neurodegeneration. Sci Transl Med 9

Kanai Y, Hediger MA (2004) The glutamate/neutral amino acid transporter family SLC1: molecular, physiological and pharmacological aspects. Pflugers Arch 447: 469–479

Kang MJ, Abdelmohsen K, Hutchison ER, Mitchell SJ, Grammatikakis I, Guo R, Noh JH, Martindale JL, Yang X, Lee EK et al (2014) HuD regulates coding and noncoding RNA to induce APP-->Abeta processing. Cell Rep 7: 1401–1409

Kato S, Sumi-Akamaru H, Fujimura H, Sakoda S, Kato M, Hirano A, Takikawa M, Ohama E (2001) Copper chaperone for superoxide dismutase co-aggregates with superoxide dismutase 1 (SOD1) in neuronal Lewy body-like hyaline inclusions: an immunohistochemical study on familial amyotrophic lateral sclerosis with SOD1 gene mutation. Acta Neuropathol 102: 233–238

Key J, Almaguer-Mederos LE, Kandi AR, Sen NE, Gispert S, Kopf G, Meierhofer D, Auburger G (2025) ATXN2L primarily interacts with NUFIP2, the absence of ATXN2L results in NUFIP2 depletion, and the ATXN2-polyQ expansion triggers NUFIP2 accumulation. Neurobiol Dis 209: 106903

Kikin O, D’Antonio L, Bagga PS (2006) QGRS Mapper: a web-based server for predicting G-quadruplexes in nucleotide sequences. Nucleic Acids Res 34: W676–682

Li M, Du Y, Zhang X, Zhou W (2024) Research advances of MAL family members in tumorigenesis and tumor progression (Review). Mol Med Rep 29

Liao JY, Yang B, Zhang YC, Wang XJ, Ye Y, Peng JW, Yang ZZ, He JH, Zhang Y, Hu K et al (2020) EuRBPDB: a comprehensive resource for annotation, functional and oncological investigation of eukaryotic RNA binding proteins (RBPs). Nucleic Acids Res 48: D307–D313

Loughlin FE, Lukavsky PJ, Kazeeva T, Reber S, Hock EM, Colombo M, Von Schroetter C, Pauli P, Clery A, Muhlemann O et al (2019) The Solution Structure of FUS Bound to RNA Reveals a Bipartite Mode of RNA Recognition with Both Sequence and Shape Specificity. Mol Cell 73: 490–504 e496

Mackenzie IR, Bigio EH, Ince PG, Geser F, Neumann M, Cairns NJ, Kwong LK, Forman MS, Ravits J, Stewart H et al (2007) Pathological TDP-43 distinguishes sporadic amyotrophic lateral sclerosis from amyotrophic lateral sclerosis with SOD1 mutations. Ann Neurol 61: 427–434

Marsh AP (2019) Molecular mechanisms of proteinopathies across neurodegenerative disease: a review. Neurol Res Pract 1: 35

Martin LJ, Liu Z, Chen K, Price AC, Pan Y, Swaby JA, Golden WC (2007) Motor neuron degeneration in amyotrophic lateral sclerosis mutant superoxide dismutase-1 transgenic mice: mechanisms of mitochondriopathy and cell death. J Comp Neurol 500: 20–46

Masrori P, Van Damme P (2020) Amyotrophic lateral sclerosis: a clinical review. Eur J Neurol 27: 1918–1929

Matsuo K, Asamitsu S, Maeda K, Suzuki H, Kawakubo K, Komiya G, Kudo K, Sakai Y, Hori K, Ikenoshita S et al (2024) RNA G-quadruplexes form scaffolds that promote neuropathological alpha-synuclein aggregation. Cell 187: 6835–6848 e6820

Mead RJ, Shan N, Reiser HJ, Marshall F, Shaw PJ (2023) Amyotrophic lateral sclerosis: a neurodegenerative disorder poised for successful therapeutic translation. Nat Rev Drug Discov 22: 185–212

Montalbano M, McAllen S, Cascio FL, Sengupta U, Garcia S, Bhatt N, Ellsworth A, Heidelman EA, Johnson OD, Doskocil S et al (2020) TDP-43 and Tau Oligomers in Alzheimer’s Disease, Amyotrophic Lateral Sclerosis, and Frontotemporal Dementia. Neurobiol Dis 146: 105130

Mori K, Weng SM, Arzberger T, May S, Rentzsch K, Kremmer E, Schmid B, Kretzschmar HA, Cruts M, Van Broeckhoven C et al (2013) The C9orf72 GGGGCC repeat is translated into aggregating dipeptide-repeat proteins in FTLD/ALS. Science 339: 1335–1338

Moussaud S, Jones DR, Moussaud-Lamodiere EL, Delenclos M, Ross OA, McLean PJ (2014) Alpha-synuclein and tau: teammates in neurodegeneration? Mol Neurodegener 9: 43

Murakami K (2014) Conformation-specific antibodies to target amyloid β oligomers and their application to immunotherapy for Alzheimer’s disease. Biosci Biotechnol Biochem 78: 1293–1305

Murakami K, Izuo N, Bitan G (2022) Aptamers targeting amyloidogenic proteins and their emerging role in neurodegenerative diseases. J Biol Chem 298: 101478

Murakami K, Ono K (2022) Interactions of amyloid coaggregates with biomolecules and its relevance to neurodegeneration. FASEB J 36: e22493

Murakami K, Sakaguchi Y, Taniwa K, Izuo N, Hanaki M, Kawase T, Hirose K, Shimizu T, Irie K (2022) Lysine-targeting inhibition of amyloid beta oligomerization by a green perilla-derived metastable chalcone in vitro and in vivo. RSC Chem Biol 3: 1380–1396

Murakami K, Nguyen THV, Nagao C, Mizuguchi K, Bitan G (2025) Lysine-targeting inhibitors of amyloidogenic protein aggregation: a promise for neurodegenerative proteinopathies. JACS Au doi: 10.1021/jacsau.5c00269

Neumann M, Sampathu DM, Kwong LK, Truax AC, Micsenyi MC, Chou TT, Bruce J, Schuck T, Grossman M, Clark CM et al (2006) Ubiquitinated TDP-43 in frontotemporal lobar degeneration and amyotrophic lateral sclerosis. Science 314: 130–133

Piovesan D, Del Conte A, Mehdiabadi M, Aspromonte MC, Blum M, Tesei G, von Bulow S, Lindorff-Larsen K, Tosatto SCE (2025) MOBIDB in 2025: integrating ensemble properties and function annotations for intrinsically disordered proteins. Nucleic Acids Res 53: D495–D503

Ramos-Campoy O, Avila-Polo R, Grau-Rivera O, Antonell A, Clarimon J, Rojas-Garcia R, Charif S, Santiago-Valera V, Hernandez I, Aguilar M et al (2018) Systematic Screening of Ubiquitin/p62 Aggregates in Cerebellar Cortex Expands the Neuropathological Phenotype of the C9orf72 Expansion Mutation. J Neuropathol Exp Neurol 77: 703–709

Roychaudhuri R, Yang M, Hoshi MM, Teplow DB (2009) Amyloid β-protein assembly and Alzheimer disease. J Biol Chem 284: 4749–4753

Sathyaseelan C, Vijayakumar V, Rathinavelan T (2021) CD-NuSS: A Web Server for the Automated Secondary Structural Characterization of the Nucleic Acids from Circular Dichroism Spectra Using Extreme Gradient Boosting Decision-Tree, Neural Network and Kohonen Algorithms. J Mol Biol 433: 166629

Selkoe DJ (2001) Alzheimer’s disease: genes, proteins, and therapy. Physiol Rev 81: 741–766

Silvestri B, Mochi M, Mawrie D, de Turris V, Colantoni A, Borhy B, Medici M, Anderson EN, Garone MG, Zammerilla CP et al (2024) HuD impairs neuromuscular junctions and induces apoptosis in human iPSC and Drosophila ALS models. Nat Commun 15: 9618

Spencer S, Dowbenko D, Cheng J, Li W, Brush J, Utzig S, Simanis V, Lasky LA (1997) PSTPIP: a tyrosine phosphorylated cleavage furrow-associated protein that is a substrate for a PEST tyrosine phosphatase. J Cell Biol 138: 845–860

Spillantini MG, Goedert M (2000) The alpha-synucleinopathies: Parkinson’s disease, dementia with Lewy bodies, and multiple system atrophy. Ann N Y Acad Sci 920: 16–27

Spires-Jones TL, Attems J, Thal DR (2017) Interactions of pathological proteins in neurodegenerative diseases. Acta Neuropathol 134: 187–205

Subramanian M, Rage F, Tabet R, Flatter E, Mandel JL, Moine H (2011) G-quadruplex RNA structure as a signal for neurite mRNA targeting. EMBO Rep 12: 697–704

Taylor JP, Brown RH, Jr., Cleveland DW (2016) Decoding ALS: from genes to mechanism. Nature 539: 197–206

Vance C, Rogelj B, Hortobagyi T, De Vos KJ, Nishimura AL, Sreedharan J, Hu X, Smith B, Ruddy D, Wright P et al (2009) Mutations in FUS, an RNA processing protein, cause familial amyotrophic lateral sclerosis type 6. Science 323: 1208–1211

Wolozin B, Ivanov P (2019) Stress granules and neurodegeneration. Nat Rev Neurosci 20: 649–666

Xie L, Zhu Y, Hurtle BT, Wright M, Robinson JL, Mauna JC, Brown EE, Ngo M, Bergmann CA, Xu J et al (2025) Context-dependent Interactors Regulate TDP-43 Dysfunction in ALS/FTLD. bioRxiv

Yabuki Y, Matsuo K, Komiya G, Kudo K, Hori K, Ikenoshita S, Kawata Y, Mizobata T, Shioda N (2024) RNA G-quadruplexes and calcium ions synergistically induce Tau phase transition in vitro. J Biol Chem 300: 107971

Yagi R, Miyazaki T, Oyoshi T (2018) G-quadruplex binding ability of TLS/FUS depends on the beta-spiral structure of the RGG domain. Nucleic Acids Res 46: 5894–5901

Zekry D, Hauw JJ, Gold G (2002) Mixed dementia: epidemiology, diagnosis, and treatment. J Am Geriatr Soc 50: 1431–1438

