## Supporting Information Table S1 for "Proteomic and *in silico* dissection of MetaAggregates in amyotrophic lateral sclerosis brains"

**Authors' Information**
ORCiD 0000-0003-3152-1784 (K.M.)
ORCiD 0000-0001-8454-6155 (K.O.)

---

[Contents]

**Table S1.** RMSD (Å) and other parameters (arbitrary units) from docking simulations assessing the structural association between ELAVL4 and ALS-associated proteins.

(A) ELAVL4 vs TDP-43
(B) ELAVL4 vs SRP9
(C) ELAVL4 vs NDUFA2
(D) ELAVL4 vs NUFIP2
(E) ELAVL4 vs AVEN
(F) ELAVL4 vs MAL2
(G) ELAVL4 vs SLC1A3
(H) ELAVL4 vs CAPS
(I) ELAVL4 vs AK3
(J) ELAVL4 vs PFDN2
(K) ELAVL4 vs STIP1
(L) ELAVL4 vs HTRA2

**Table S1.** RMSD (Å) and other parameters (arbitrary units) from docking simulations assessing the structural association between ELAVL4 and ALS-associated proteins.

(A) ELAVL4 vs TDP-43

| Structures | HADDOCK score | RMSD from the overall lowest-energy structure | Z-Score | Van der Waals energy |
|---|---|---|---|---|
| 1FXL_6T4B_Cluster2.pdb | -128.2 +/- 5.9 | 0.6 +/- 0.4 | -2.3 | -50.8 +/- 9.3 |
| 1FXL_6T4B_Cluster8.pdb | -106.9 +/- 11.0 | 6.4 +/- 0.3 | -0.8 | -45.7 +/- 10.5 |
| 1FXL_6T4B_Cluster3.pdb | -101.3 +/- 6.1 | 5.3 +/- 0.5 | -0.4 | -44.4 +/- 10.2 |
| 1FXL_6T4B_Cluster5.pdb | -98.7 +/- 9.3 | 9.6 +/- 0.2 | -0.2 | -61.9 +/- 6.9 |
| 1FXL_6T4B_Cluster4.pdb | -96.2 +/- 4.4 | 8.3 +/- 0.4 | 0 | -50.9 +/- 6.2 |
| 1FXL_6T4B_Cluster7.pdb | -95.6 +/- 16.0 | 9.7 +/- 0.1 | 0.1 | -52.5 +/- 16.4 |
| 1FXL_6T4B_Cluster1.pdb | -92.9 +/- 5.4 | 6.1 +/- 0.3 | 0.3 | -45.0 +/- 4.5 |
| 1FXL_6T4B_Cluster13.pdb | -85.8 +/- 14.2 | 8.7 +/- 0.2 | 0.8 | -52.2 +/- 8.6 |
| 1FXL_6T4B_Cluster6.pdb | -82.9 +/- 3.6 | 8.1 +/- 0.3 | 1 | -40.4 +/- 2.4 |
| 1FXL_6T4B_Cluster11.pdb | -76.7 +/- 30.9 | 5.9 +/- 0.3 | 1.5 | -38.8 +/- 14.1 |

### (B) ELAVL4 vs SRP9

| Structures | HADDOCK score | RMSD from the overall lowest-energy structure | Z-Score | Van der Waals energy |
|---|---|---|---|---|
| 1FXL_1914_Cluster4.pdb | -122.3 +/- 14.9 | 1.2 +/- 0.5 | -1.5 | -61.1 +/- 5.2 |
| 1FXL_1914_Cluster2.pdb | -111.4 +/- 3.9 | 9.3 +/- 0.7 | -1 | -66.1 +/- 4.0 |
| 1FXL_1914_Cluster9.pdb | -106.2 +/- 3.0 | 1.4 +/- 1.0 | -2.1 | -64.0 +/- 10.7 |
| 1FXL_1914_Cluster1.pdb | -99.1 +/- 4.0 | 6.6 +/- 0.6 | -0.4 | -62.8 +/- 2.9 |
| 1FXL_1914_Cluster7.pdb | -96.9 +/- 4.6 | 17.8 +/- 0.1 | -1.5 | -69.8 +/- 5.5 |
| 1FXL_1914_Cluster5.pdb | -93.7 +/- 3.9 | 16.6 +/- 0.5 | -0.2 | -58.8 +/- 2.8 |
| 1FXL_1914_Cluster8.pdb | -74.9 +/- 3.8 | 17.6 +/- 0.7 | -0.2 | -58.9 +/- 6.8 |
| 1FXL_1914_Cluster3.pdb | -71.4 +/- 7.2 | 4.5 +/- 0.7 | 0.8 | -40.7 +/- 4.6 |
| 1FXL_1914_Cluster6.pdb | -70.6 +/- 7.6 | 6.1 +/- 0.3 | 0.9 | -33.4 +/- 1.5 |
| 1FXL_1914_Cluster10.pdb | -69.2 +/- 10.7 | 9.2 +/- 0.8 | 0.2 | -58.9 +/- 7.5 |

(C) ELAVL4 vs NDUFA2

| Structures | HADDOCK score | RMSD from the overall lowest-energy structure | Z-Score | Van der Waals energy |
|---|---|---|---|---|
| 1FXL_1S3A_Cluster4.pdb | -122.2 +/- 3.8 | 0.4 +/- 0.3 | -2.2 | -56.4 +/- 6.9 |
| 1FXL_1S3A_Cluster1.pdb | -101.9 +/- 1.4 | 11.8 +/- 0.2 | -0.9 | -54.5 +/- 4.5 |
| 1FXL_1S3A_Cluster8.pdb | -100.1 +/- 9.2 | 4.9 +/- 0.0 | -0.8 | -74.1 +/- 6.4 |
| 1FXL_1S3A_Cluster5.pdb | -88.4 +/- 7.4 | 6.8 +/- 0.3 | 0 | -62.8 +/- 6.1 |
| 1FXL_1S3A_Cluster6.pdb | -87.4 +/- 9.0 | 3.6 +/- 0.1 | 0 | -50.5 +/- 5.6 |
| 1FXL_1S3A_Cluster13.pdb | -83.8 +/- 6.2 | 10.7 +/- 0.2 | 0.2 | -50.4 +/- 5.5 |
| 1FXL_1S3A_Cluster2.pdb | -81.1 +/- 9.7 | 6.4 +/- 0.3 | 0.4 | -42.2 +/- 3.6 |
| 1FXL_1S3A_Cluster3.pdb | -77.3 +/- 8.2 | 11.7 +/- 0.1 | 0.7 | -60.6 +/- 4.3 |
| 1FXL_1S3A_Cluster12.pdb | -67.8 +/- 3.6 | 5.4 +/- 0.3 | 1.2 | -39.3 +/- 4.1 |
| 1FXL_1S3A_Cluster9.pdb | -67.1 +/- 14.2 | 9.6 +/- 0.4 | 1.3 | -43.4 +/- 8.8 |

(D) ELAVL4 vs NUFIP2

| Structures | HADDOCK score | RMSD from the overall lowest-energy structure | Z-Score | Van der Waals energy |
|---|---|---|---|---|
| 1FXL_AF-Q7Z417-F1_Cluster1.pdb | -117.9 +/- 11.1 | 9.5 +/- 0.1 | -2.4 | -75.7 +/- 7.8 |
| 1FXL_AF-Q7Z417-F1_Cluster9.pdb | -68.5 +/- 8.8 | 14.5 +/- 0.2 | -0.7 | -47.2 +/- 8.0 |
| 1FXL_AF-Q7Z417-F1_Cluster2.pdb | -58.7 +/- 12.7 | 14.6 +/- 0.2 | -0.4 | -51.4 +/- 6.9 |
| 1FXL_AF-Q7Z417-F1_Cluster10.pdb | -47.7 +/- 16.1 | 9.3 +/- 0.1 | 0 | -45.3 +/- 6.8 |
| 1FXL_AF-Q7Z417-F1_Cluster6.pdb | -46.0 +/- 8.9 | 14.8 +/- 0.0 | 0 | -33.4 +/- 7.7 |
| 1FXL_AF-Q7Z417-F1_Cluster5.pdb | -41.7 +/- 13.9 | 9.3 +/- 0.0 | 0.2 | -51.1 +/- 9.8 |
| 1FXL_AF-Q7Z417-F1_Cluster4.pdb | -33.4 +/- 6.8 | 8.0 +/- 0.2 | 0.4 | -35.9 +/- 11.8 |
| 1FXL_AF-Q7Z417-F1_Cluster7.pdb | -22.4 +/- 10.7 | 14.7 +/- 0.1 | 0.8 | -37.6 +/- 4.8 |
| 1FXL_AF-Q7Z417-F1_Cluster8.pdb | -21.6 +/- 0.5 | 10.6 +/- 0.2 | 0.8 | -17.1 +/- 4.0 |
| 1FXL_AF-Q7Z417-F1_Cluster3.pdb | -6.1 +/- 8.0 | 9.7 +/- 0.0 | 1.4 | -27.9 +/- 6.2 |

(E) ELAVL4 vs AVEN

| Structures | HADDOCK score | RMSD from the overall lowest-energy structure | Z-Score | Van der Waals energy |
|---|---|---|---|---|
| 1FXL_AF-Q9NQS1-F1_Cluster7.pdb | -158.0 +/- 17.1 | 1.8 +/- 1.2 | -2 | -60.5 +/- 8.8 |
| 1FXL_AF-Q9NQS1-F1_Cluster10.pdb | -124.8 +/- 13.9 | 16.7 +/- 0.3 | -0.8 | -64.8 +/- 9.5 |
| 1FXL_AF-Q9NQS1-F1_Cluster4.pdb | -116.5 +/- 8.4 | 18.1 +/- 0.3 | -0.5 | -67.7 +/- 7.4 |
| 1FXL_AF-Q9NQS1-F1_Cluster5.pdb | -113.4 +/- 10.7 | 18.2 +/- 0.1 | -0.4 | -84.3 +/- 4.0 |
| 1FXL_AF-Q9NQS1-F1_Cluster6.pdb | -105.5 +/- 3.2 | 10.9 +/- 0.3 | -0.1 | -69.2 +/- 9.8 |
| 1FXL_AF-Q9NQS1-F1_Cluster2.pdb | -105.2 +/- 4.8 | 6.7 +/- 0.2 | -0.1 | -51.3 +/- 4.9 |
| 1FXL_AF-Q9NQS1-F1_Cluster1.pdb | -92.7 +/- 2.6 | 18.2 +/- 0.3 | 0.4 | -58.8 +/- 9.7 |
| 1FXL_AF-Q9NQS1-F1_Cluster3.pdb | -87.2 +/- 10.4 | 17.9 +/- 0.1 | 0.6 | -65.0 +/- 10.5 |
| 1FXL_AF-Q9NQS1-F1_Cluster13.pdb | -65.6 +/- 10.5 | 7.9 +/- 0.9 | 1.3 | -47.4 +/- 6.1 |
| 1FXL_AF-Q9NQS1-F1_Cluster11.pdb | -56.6 +/- 22.3 | 17.0 +/- 0.2 | 1.7 | -46.4 +/- 12.7 |

(F) ELAVL4 vs MAL2

| Structures | HADDOCK score | RMSD from the overall lowest-energy structure | Z-Score | Van der Waals energy |
|---|---|---|---|---|
| 1FXL_AF-Q969L2-F1_Cluster1.pdb | -123.3 +/- 0.9 | 23.1 +/- 0.1 | -1.5 | -92.0 +/- 4.6 |
| 1FXL_AF-Q969L2-F1_Cluster7.pdb | -119.3 +/- 8.0 | 10.5 +/- 0.2 | -1.3 | -69.9 +/- 9.9 |
| 1FXL_AF-Q969L2-F1_Cluster11.pdb | -110.4 +/- 26.3 | 1.3 +/- 0.8 | -0.7 | -63.8 +/- 11.6 |
| 1FXL_AF-Q969L2-F1_Cluster4.pdb | -105.6 +/- 12.0 | 22.4 +/- 0.5 | -0.4 | -65.7 +/- 10.2 |
| 1FXL_AF-Q969L2-F1_Cluster2.pdb | -104.8 +/- 9.3 | 10.6 +/- 0.2 | -0.4 | -75.5 +/- 9.0 |
| 1FXL_AF-Q969L2-F1_Cluster3.pdb | -97.8 +/- 2.7 | 24.4 +/- 0.1 | 0 | -48.7 +/- 4.9 |
| 1FXL_AF-Q969L2-F1_Cluster10.pdb | -91.5 +/- 22.4 | 5.0 +/- 0.2 | 0.4 | -56.9 +/- 12.0 |
| 1FXL_AF-Q969L2-F1_Cluster5.pdb | -77.7 +/- 8.5 | 12.2 +/- 0.5 | 1.2 | -63.5 +/- 12.1 |
| 1FXL_AF-Q969L2-F1_Cluster6.pdb | -76.2 +/- 16.9 | 23.3 +/- 0.1 | 1.3 | -66.2 +/- 6.9 |
| 1FXL_AF-Q969L2-F1_Cluster8.pdb | -73.7 +/- 20.9 | 23.4 +/- 0.1 | 1.4 | -57.6 +/- 9.3 |

### (G) ELAVL4 vs SLC1A3

| Structures | HADDOCK score | RMSD from the overall lowest-energy structure | Z-Score | Van der Waals energy |
|---|---|---|---|---|
| 1FXL_5LLU_Cluster2.pdb | -110.4 +/- 4.1 | 0.6 +/- 0.6 | -1.1 | -57.4 +/- 2.4 |
| 1FXL_5LLU_Cluster3.pdb | -108.1 +/- 5.9 | 16.5 +/- 0.0 | -1.7 | -51.6 +/- 4.2 |
| 1FXL_5LLU_Cluster22.pdb | -102.5 +/- 19.3 | 5.1 +/- 0.6 | -1 | -59.5 +/- 10.0 |
| 1FXL_5LLU_Cluster1.pdb | -100.6 +/- 2.6 | 16.8 +/- 0.4 | -0.8 | -53.8 +/- 2.5 |
| 1FXL_5LLU_Cluster6.pdb | -98.2 +/- 16.0 | 18.6 +/- 0.0 | -0.5 | -65.7 +/- 10.8 |
| 1FXL_5LLU_Cluster5.pdb | -96.7 +/- 2.4 | 2.4 +/- 0.3 | -0.3 | -44.1 +/- 6.0 |
| 1FXL_5LLU_Cluster10.pdb | -93.9 +/- 5.0 | 11.3 +/- 0.3 | 0.1 | -40.1 +/- 5.4 |
| 1FXL_5LLU_Cluster21.pdb | -88.9 +/- 11.8 | 16.4 +/- 0.2 | 0.7 | -53.2 +/- 9.6 |
| 1FXL_5LLU_Cluster14.pdb | -88.1 +/- 9.7 | 16.3 +/- 0.2 | 0.8 | -45.3 +/- 8.7 |
| 1FXL_5LLU_Cluster4.pdb | -83.8 +/- 3.4 | 14.8 +/- 0.5 | 1.4 | -51.2 +/- 7.6 |

(H) ELAVL4 vs CAPS

| Structures | HADDOCK score | RMSD from the overall lowest-energy structure | Z-Score | Van der Waals energy |
|---|---|---|---|---|
| 1FXL_3E3R_Cluster5.pdb | -111.9 +/- 11.6 | 0.9 +/- 0.5 | -2.2 | -76.0 +/- 11.9 |
| 1FXL_3E3R_Cluster11.pdb | -94.3 +/- 7.3 | 23.3 +/- 0.1 | -1.2 | -47.9 +/- 7.8 |
| 1FXL_3E3R_Cluster9.pdb | -75.2 +/- 14.7 | 23.6 +/- 0.5 | -0.2 | -34.0 +/- 7.0 |
| 1FXL_3E3R_Cluster13.pdb | -71.0 +/- 4.5 | 22.2 +/- 0.3 | 0 | -63.8 +/- 9.3 |
| 1FXL_3E3R_Cluster10.pdb | -70.9 +/- 4.6 | 10.6 +/- 0.5 | 0.1 | -34.9 +/- 3.2 |
| 1FXL_3E3R_Cluster2.pdb | -70.3 +/- 5.7 | 18.0 +/- 0.3 | 0.1 | -53.0 +/- 5.2 |
| 1FXL_3E3R_Cluster4.pdb | -64.5 +/- 5.6 | 11.4 +/- 0.4 | 0.4 | -52.9 +/- 2.1 |
| 1FXL_3E3R_Cluster8.pdb | -60.9 +/- 15.0 | 23.3 +/- 0.3 | 0.6 | -37.5 +/- 9.0 |
| 1FXL_3E3R_Cluster1.pdb | -49.5 +/- 4.2 | 22.1 +/- 1.0 | 1.2 | -52.7 +/- 2.9 |
| 1FXL_3E3R_Cluster3.pdb | -49.2 +/- 6.6 | 10.6 +/- 0.2 | 1.2 | -48.8 +/- 8.2 |

(I) ELAVL4 vs AK3

| Structures | HADDOCK score | RMSD from the overall lowest-energy structure | Z-Score | Van der Waals energy |
|---|---|---|---|---|
| 1FXL_6ZJB_Cluster1.pdb | -106.5 +/- 1.5 | 1.2 +/- 0.8 | -2.6 | -79.0 +/- 9.6 |
| 1FXL_6ZJB_Cluster8.pdb | -79.6 +/- 13.0 | 18.9 +/- 0.1 | -0.7 | -69.6 +/- 8.3 |
| 1FXL_6ZJB_Cluster14.pdb | -71.4 +/- 13.5 | 2.5 +/- 0.3 | -0.2 | -61.0 +/- 12.9 |
| 1FXL_6ZJB_Cluster2.pdb | -67.1 +/- 20.6 | 20.3 +/- 0.2 | 0.1 | -53.5 +/- 5.7 |
| 1FXL_6ZJB_Cluster10.pdb | -66.2 +/- 8.7 | 3.3 +/- 0.6 | 0.1 | -66.1 +/- 14.8 |
| 1FXL_6ZJB_Cluster3.pdb | -64.5 +/- 7.5 | 3.1 +/- 0.5 | 0.3 | -63.8 +/- 9.8 |
| 1FXL_6ZJB_Cluster12.pdb | -63.0 +/- 22.8 | 5.3 +/- 0.7 | 0.4 | -53.6 +/- 16.1 |
| 1FXL_6ZJB_Cluster4.pdb | -59.2 +/- 11.1 | 19.5 +/- 0.5 | 0.6 | -58.6 +/- 4.1 |
| 1FXL_6ZJB_Cluster13.pdb | -57.5 +/- 24.4 | 10.6 +/- 0.1 | 0.7 | -32.2 +/- 15.8 |
| 1FXL_6ZJB_Cluster5.pdb | -49.2 +/- 6.9 | 20.5 +/- 1.0 | 1.3 | -61.0 +/- 5.3 |

(J) ELAVL4 vs PFDN2

| Structures | HADDOCK score | RMSD from the overall lowest-energy structure | Z-Score | Van der Waals energy |
|---|---|---|---|---|
| 1FXL_6NR8_Cluster1.pdb | -107.5 +/- 8.1 | 12.0 +/- 0.2 | -1.7 | -47.5 +/- 5.5 |
| 1FXL_6NR8_Cluster8.pdb | -100.0 +/- 20.6 | 0.9 +/- 0.6 | -1.3 | -59.6 +/- 10.8 |
| 1FXL_6NR8_Cluster2.pdb | -97.4 +/- 4.2 | 14.5 +/- 0.4 | -1.1 | -43.5 +/- 6.2 |
| 1FXL_6NR8_Cluster11.pdb | -84.8 +/- 17.0 | 13.3 +/- 1.1 | -0.5 | -40.2 +/- 7.0 |
| 1FXL_6NR8_Cluster7.pdb | -70.4 +/- 6.1 | 3.0 +/- 0.2 | 0.2 | -37.5 +/- 7.0 |
| 1FXL_6NR8_Cluster6.pdb | -66.1 +/- 10.3 | 12.9 +/- 0.6 | 0.5 | -33.6 +/- 6.6 |
| 1FXL_6NR8_Cluster4.pdb | -63.4 +/- 10.1 | 16.3 +/- 0.8 | 0.6 | -31.9 +/- 7.8 |
| 1FXL_6NR8_Cluster9.pdb | -55.2 +/- 13.2 | 12.5 +/- 0.2 | 1 | -34.2 +/- 5.6 |
| 1FXL_6NR8_Cluster10.pdb | -54.6 +/- 4.7 | 10.7 +/- 0.1 | 1 | -25.1 +/- 5.3 |
| 1FXL_6NR8_Cluster3.pdb | -52.1 +/- 4.4 | 20.9 +/- 0.3 | 1.2 | -33.2 +/- 4.4 |

(K) ELAVL4 vs STIP1

| Structures | HADDOCK score | RMSD from the overall lowest-energy structure | Z-Score | Van der Waals energy |
|---|---|---|---|---|
| 1FXL_3Q47_Cluster10.pdb | -106.5 +/- 12.1 | 0.6 +/- 0.4 | -1.5 | -66.8 +/- 7.9 |
| 1FXL_3Q47_Cluster1.pdb | -102.3 +/- 5.1 | 2.1 +/- 0.7 | -1.3 | -69.3 +/- 5.6 |
| 1FXL_3Q47_Cluster2.pdb | -100.9 +/- 5.2 | 15.7 +/- 0.2 | -1.3 | -48.3 +/- 6.1 |
| 1FXL_3Q47_Cluster4.pdb | -78.2 +/- 4.3 | 17.2 +/- 0.2 | -0.2 | -66.0 +/- 5.0 |
| 1FXL_3Q47_Cluster3.pdb | -74.1 +/- 8.3 | 5.1 +/- 0.8 | 0 | -53.9 +/- 7.8 |
| 1FXL_3Q47_Cluster6.pdb | -67.7 +/- 9.2 | 11.1 +/- 0.3 | 0.2 | -53.6 +/- 3.7 |
| 1FXL_3Q47_Cluster9.pdb | -55.2 +/- 2.8 | 8.9 +/- 0.1 | 0.8 | -41.8 +/- 6.7 |
| 1FXL_3Q47_Cluster8.pdb | -53.2 +/- 8.4 | 10.7 +/- 1.1 | 0.9 | -57.1 +/- 4.0 |
| 1FXL_3Q47_Cluster5.pdb | -50.2 +/- 2.3 | 4.4 +/- 0.6 | 1 | -38.1 +/- 3.6 |
| 1FXL_3Q47_Cluster7.pdb | -43.1 +/- 9.7 | 12.7 +/- 0.1 | 1.4 | -43.9 +/- 4.2 |

(L) ELAVL4 vs HTRA2

| Structures | HADDOCK score | RMSD from the overall lowest-energy structure | Z-Score | Van der Waals energy |
|---|---|---|---|---|
| 1FXL_1LCY_Cluster3.pdb | -115.5 +/- 11.7 | 2.1 +/- 1.7 | -2.5 | -89.1 +/- 9.6 |
| 1FXL_1LCY_Cluster8.pdb | -92.3 +/- 4.1 | 10.8 +/- 0.3 | -0.8 | -66.2 +/- 8.4 |
| 1FXL_1LCY_Cluster7.pdb | -89.7 +/- 9.4 | 13.5 +/- 0.2 | -0.6 | -73.3 +/- 4.9 |
| 1FXL_1LCY_Cluster13.pdb | -84.3 +/- 22.0 | 17.2 +/- 0.5 | -0.2 | -43.6 +/- 16.2 |
| 1FXL_1LCY_Cluster1.pdb | -78.5 +/- 3.5 | 11.4 +/- 1.1 | 0.3 | -51.1 +/- 5.6 |
| 1FXL_1LCY_Cluster11.pdb | -75.3 +/- 8.9 | 14.6 +/- 0.4 | 0.5 | -59.9 +/- 9.9 |
| 1FXL_1LCY_Cluster2.pdb | -72.2 +/- 3.0 | 13.2 +/- 0.9 | 0.7 | -45.1 +/- 11.7 |
| 1FXL_1LCY_Cluster5.pdb | -72.0 +/- 4.5 | 13.7 +/- 0.4 | 0.7 | -56.2 +/- 8.2 |
| 1FXL_1LCY_Cluster4.pdb | -69.7 +/- 7.9 | 18.7 +/- 0.0 | 0.9 | -51.7 +/- 7.2 |
| 1FXL_1LCY_Cluster6.pdb | -31.5 +/- 23.3 | 23.4 +/- 0.3 | 0.8 | -50.8 +/- 9.2 |
